# Epigenetic Control of TERRA by FTSJ3 is Critical for Telomerase-Driven Cancers

**DOI:** 10.1101/2025.01.26.634101

**Authors:** Jared D.W. Price, Frederick S. Vizeacoumar, Omar Abuhussein, Vincent Maranda, Yue Zhang, Hironori Adachi, Liliia Kyrylenko, Aline Rangel-Pozzo, He Dong, Li Hui Gong, Prachi Walke, Ashtalakshmi Ganapathysamy, Connor Denomy, Tanya Freywald, Renuka Dahiya, Hussain Elhasasna, Anjali Saxena, Jeff P. Vizeacoumar, Hardikkumar Patel, Karthic Rajamanickam, Kathryn Nguyen, Diego M. de Oliveira, Mary Lazell-Wright, Alain Morejon Morales, Aanchal Aggarwal, Jia Lin Xu, Nezeka Alli, Erika P. Munhoz, Peng Gao, Jayme Salsman, Dinesh Dahiya, Cristina Gonzalez-Lopez, Patricia Thibault, Michael Levin, Graham Dellaire, Nicholas Jette, Gary Groot, Anand Krishnan, Shahid Ahmed, Christopher Eskiw, Khaled H. Barakat, Yuliang Wu, Ronald A. DePinho, Sabine Mai, Yi-Tao Yu, Judy M.Y. Wong, Andrew Freywald, Franco J. Vizeacoumar

## Abstract

Telomerase reverse transcriptase (hTERT) overexpression, a hallmark of most cancers, drives tumorigenesis by enabling limitless replicative potential. Direct targeting of hTERT is challenging, necessitating alternative strategies. Through genome-wide synthetic dosage lethality (SDL) screening in cancer models, including patient-derived organoids, we identify FTSJ3, an RNA 2’-O-methyltransferase, as a critical vulnerability in hTERT-overexpressing cells. FTSJ3 methylates telomeric repeat-containing RNA (TERRA), a modification essential for recruiting SUV39H1 to telomeric ends to mediate H3K9 trimethylation and establish stable heterochromatin. Loss of FTSJ3 disrupts this cascade, impairing H3K9 trimethylation, HP1-alpha recruitment, and telomeric heterochromatin maintenance. Notably, this reveals an unexpected dependency on TERRA methylation for telomeric heterochromatin stability in hTERT-driven cancers. Non-malignant cells, lacking telomerase activity and *de novo* telomere repeat synthesis, are unaffected by FTSJ3 suppression. Our findings establish the FTSJ3/TERRA/SUV39H1 axis as a critical mechanism supporting telomeric heterochromatin stability in hTERT-driven cancers. This telomere-directed epigenetic strategy provides a robust framework for translational therapeutic innovation.

## Introduction

Cancer heterogeneity poses a significant medical challenge, necessitating therapies tailored not only to individual patients but also to the unique cancer clones that exist within diverse microenvironmental contexts^1^. This complexity makes the development of effective treatments particularly challenging, as variations in genetic profiles and microenvironmental interactions can influence therapy response. Despite this diversity, nearly all cancer cells rely on telomeres to maintain genomic stability^2^. Telomere maintenance, predominantly mediated by telomerase, is essential for tumor growth and viability of cancer cells. Telomerase is a ribonucleoprotein complex composed of the catalytic subunit hTERT and the RNA component hTERC. As adult somatic cells have low or undetectable levels of hTERT^3^, direct telomerase targeting has previously been expected to spare the normal cells^2^. While several telomerase targeting strategies have been developed^4^, Imetelstat is the only drug to show significant clinical benefit in heavily transfusion-dependent, lower-risk myelodysplastic syndrome (MDS) patients^5^. It is a 13-mer oligonucleotide with a lipid moiety that specifically binds to the RNA template of human telomerase. This represents a notable achievement for a niche condition like MDS, but its impact remains limited, as it is currently approved only for this specific patient population^6^.

A major limitation of direct telomerase inhibition is the slow rate of telomere shortening, which necessitates an extended lag period before critical telomere attrition is achieved to induce cell death. This requires sustained treatment over several months to elicit therapeutically meaningful tumor reduction, providing cancer cells with sufficient time to activate resistance mechanisms, such as the alternative lengthening of telomeres (ALT) pathway, which maintains telomeres through homologous recombination^7^. These challenges underscore the need for innovative strategies that effectively exploit telomerase dependencies while circumventing resistance.

To address these limitations, we employed a strategy that shifts focus from directly targeting telomerase activity to identifying genetic dependencies via synthetic dosage lethality (SDL) interactions. This approach seeks to identify genes whose inhibition selectively induces lethality in hTERT^positive^ cells, while sparing normal, hTERT^-low/negative^ cells^8^ **(Fig. 1A)**. By utilizing genome-wide CRISPR/Cas9 and shRNA screens, we aimed to uncover SDL interactions of hTERT. Through extensive validation using both arrayed *in vitro* and pooled *in vivo* CRISPR/Cas9 screens across multiple cell lines, tumor xenografts, and patient-derived organoids, we identified FTSJ3, an RNA 2’-O-methyltransferase, as a prominent SDL partner of hTERT.

**Figure 1:**
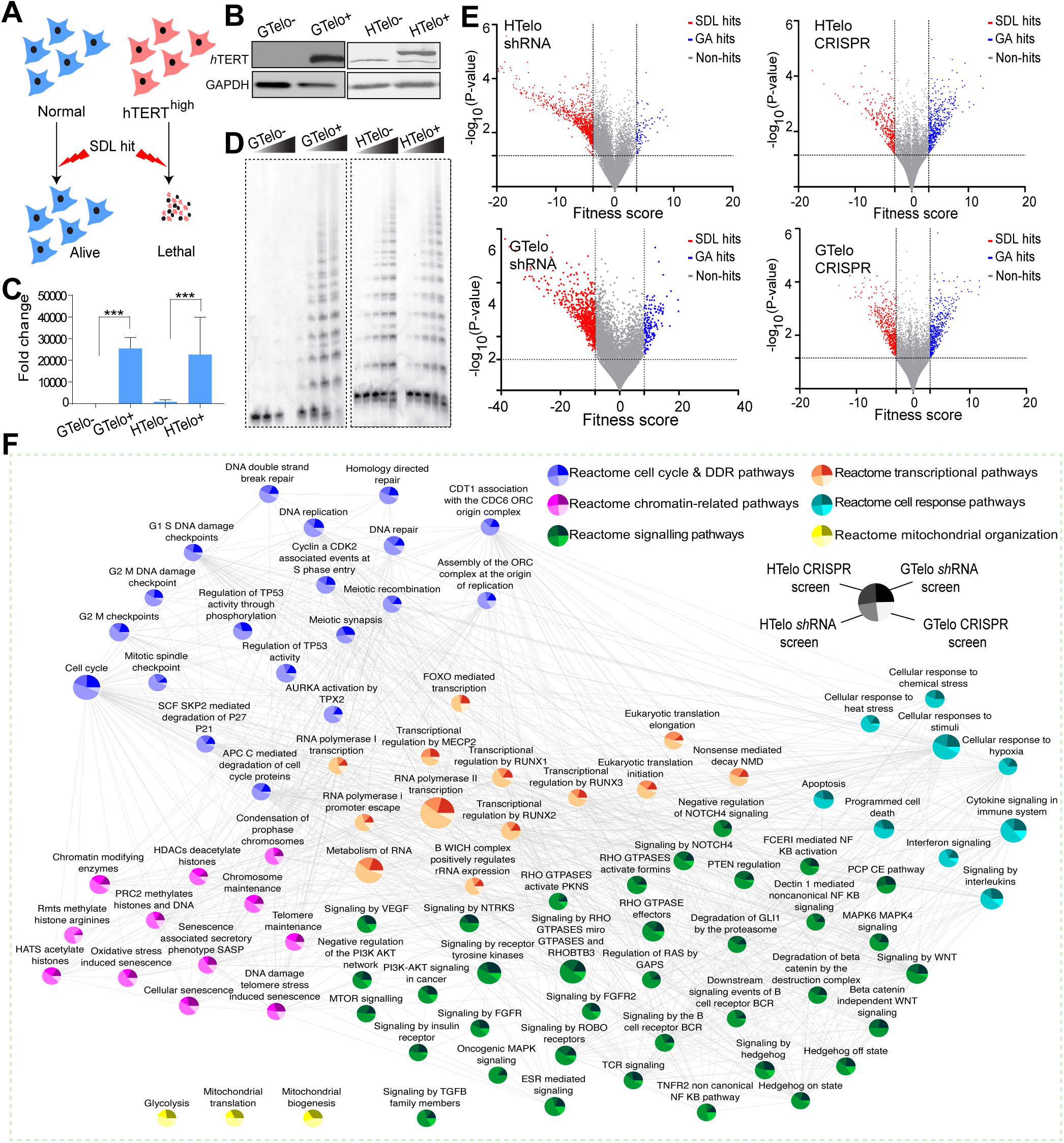
Genome-wide screen for identifying SDL interactions of hTERT. **(A)** Schematic representation depicting the concept of SDL interactions. **(B)** Western blot showing hTERT protein levels in two isogenic models. The left panel shows that the transformed fibroblasts GM00847 are hTERT negative (GTelo-) and that the derived isogenic cell line expresses hTERT (GTelo+). The right panel shows hTERT^low^ HT1080 cells (HTelo-) and the derived hTERT^high^ isogenic cell line (HTelo+). The lower band represents endogenous hTERT, and the upper band represents GFP-tagged hTERT. Western blotting with anti-GAPDH was used as a loading control. **(C)** Relative expression levels of hTERT determined by qRT‒PCR in isogenic models. Student’s t-tests; ****P<0.0001 and *P<0.05 **(D)** Enzyme activity of telomerase in the indicated isogenic models determined using a telomeric repeat amplification protocol (TRAP). Note that GTelo-cells are an ALT model and do not exhibit any telomerase activity. In HTelo+, even with forced expression of hTERT, telomerase activity remains unchanged. **(E)** Volcano plots of the genome-wide pooled screens in two isogenic models generated by two different screening approaches, i.e., shRNA screening and CRISPR/Cas9 screening. The x-axis represents the fitness score that identifies the SDL hits on the left (red dots) and those that provide a growth advantage (GA hits) on the right (blue dots). The y-axis shows the p-value significance of these hits. For statistical analysis, a t-test in conjunction with a permutation test was used to calculate the p values. **(F)** Reactome pathway enrichment analyses of all four screens. GSEA was performed, and genes with p≤0.005 and FDR≤0.1 were used to select enriched pathways. The size of the nodes represents the number of genes in each node. Each node is divided into four colors representing the percentage of genes identified in each screen.

Our findings reveal that FTSJ3 is essential for maintaining heterochromatin in hTERT^positive^ cells, specifically through methylation of telomeric repeat-containing RNA (TERRA). The loss of FTSJ3 results in a more relaxed and less stable telomeric heterochromatin structure due to inefficient loading of the histone methyltransferase SUV39H1 at the telomeric ends, as we found FTSJ3-mediated regulation of TERRA to be essential for the recruitment of SUV39H1. Given that continuous hTERT-dependent telomere synthesis and associated heterochromatinization are critical for cancer cell viability^9,10^, our results explain why FTSJ3 depletion induces a rapid onset of genomic instability, specifically in hTERT^positive^ cancer cells, while sparing normal cells that do not actively synthesize telomeric repeats. Since hTERT activity is essential in almost all cancer cells, our findings introduce a transformative approach to overcoming cancer cell heterogeneity by targeting the SDL interactions of hTERT. This strategy also capitalizes on the ubiquitous overexpression of hTERT in all cancer types, enabling the selective eradication of hTERT-dependent malignancies.

## Results

### Genome-wide SDL screens revealed genes functionally related to hTERT

To identify synthetic dosage lethality (SDL) interactions of hTERT, we employed two isogenic pairs of cell lines to reveal genes essential for the viability of hTERT^positive^ cells, using isogenic parental hTERT^low/negative^ control cells for comparison **(Fig. 1B)**. The first model involved SV40-transformed, hTERT^negative^ human fibroblast GM00847 cells (GTelo-) and an isogenic hTERT^positive^ cell line (GTelo+). Notably, GM00847 cells maintain telomeres via the telomerase-independent alternative lengthening of telomeres (ALT) mechanism^11^. Consistent with prior reports^12,13^, hTERT overexpression did not affect ALT activity, as indicated by the number of promyelocytic leukemia (PML) bodies **(Fig. S1A),** suggesting telomerase-dependent telomere synthesis and the ALT recombination mechanism could exist in co-dominantly. To account for cell line-specific effects, we also used a second model, hTERT^low^ HT1080 fibrosarcoma cells to generate N-terminally tagged GFP-hTERT^positive^ cells (referred to as HTelo- and HTelo+, respectively; **Fig. 1B**). Both models exhibit high hTERT expression **(Fig. 1C).** By using two different cell types, we enhance the robustness of our findings across diverse cellular contexts. Further characterization using the telomeric repeat amplification protocol (TRAP) demonstrated that hTERT expression reconstitutes telomerase activity in GTelo+ cells, while it remains unaltered in the HTelo-/HTelo+ model **(Fig. 1D)**. These models represent complementary mechanisms of telomere maintenance: GTelo cells employ the ALT mechanism, and the expression of hTERT allows these cells to engage two active telomere maintenance systems in concert. In contrast, HT1080 cells naturally utilize telomerase for telomere maintenance. This distinction enables us to capture both hTERT-specific effects and those associated with telomere maintenance in general.

Next, we conducted four genome-wide pooled CRISPR/Cas9 and shRNA screens, across the two isogenic pairs **(Fig. 1E**; **Fig. S1B and C)**. The quality of the screens was confirmed by high reproducibility across replicates (Pearson r > 0.89; **Fig. S1D**). To reduce false positives in the CRISPR/Cas9 screens, we employed CRISPRcleanR^14^ to account for gene dropout from copy number-amplified regions. We calculated the weighted differential cumulative change (WDC) to measure gene dropout between hTERT^positive^ and hTERT^low^ populations, using at least two sgRNAs or shRNAs per gene^15^. Despite the limited overlap between shRNA and CRISPR/Cas9 hits **(Fig. S1E; Table S1)**, pathway analysis revealed significant enrichment of both canonical and non-canonical hTERT functions (FDR < 0.05) **(Fig. 1F**, **Table S2)**. Identifying genes associated with non-canonical roles of hTERT, underscores their potential relevance to tumor development.

Confirming the validity of our approach, Reactome pathway enrichment analysis highlighted previously described associations between hTERT and the NF-κB signaling^16^, Wnt signaling pathway^17,18^, and Rho GTPases^19^, which exhibited SDL interactions **(Fig. 1F)**. Moreover, several chromatin-remodeling and DNA damage response pathways were identified, consistent with hTERT’s role in telomere maintenance. Mitochondrial pathways were also significantly enriched (FDR < 0.05; **Fig. S2A, Table S3**), supporting previous findings that anti-telomerase therapies induce mitochondrial adaptations^20^.

To further increase confidence in our screen findings, we integrated drug response data from the CancerRXgene database^21^, identifying nine chemical inhibitors that target SDL hits and preferentially suppressed hTERT^positive^ but not hTERT^low^ cells (p < 0.05; **Fig. S2B**). These chemical genetics data provide independent cross-validation of our screens and highlight their potential to uncover therapeutically relevant cancer targets.

### Validation of SDL hits identified FTSJ3 as a critical SDL target of hTERT

To increase the robustness of our findings, we employed five distinct unbiased strategies to prioritize the most likely SDL hits **(Fig. 2A)**. First, we identified 35 SDL hits co-upregulated with hTERT across 33 cancers using The Cancer Genome Atlas (TCGA) data **(Fig. S3A)**. Second, using gene expression data from two independent studies, we prioritized 21 hTERT-associated genes^20,22^ **(Fig. 2A)**. Third, we analyzed essentiality scores from DepMap^23^, identifying 14 potential SDL hits preferentially essential in hTERT^positive^ cell lines compared to hTERT^low^ or ALT cell lines (p < 0.001) **(Fig. S3B)**. Fourth, 16 clinically relevant SDL hits were prioritized by Kaplan-Meier survival analysis, distinguishing patients with ‘naturally occurring’ hTERT SDL interactions from those without (log-rank p < 0.01; **Fig. S3C**). Finally, we identified 10 SDL hits that appeared in at least three screens or ranked within the top 200 hits in any two CRISPR/Cas9 or shRNA screens. In total, 119 hits were prioritized, 21 of which failed in the cloning of sgRNAs, and we focused on validating 98 hits **(Fig. 2A**; **Table S4)**.

**Figure 2:**
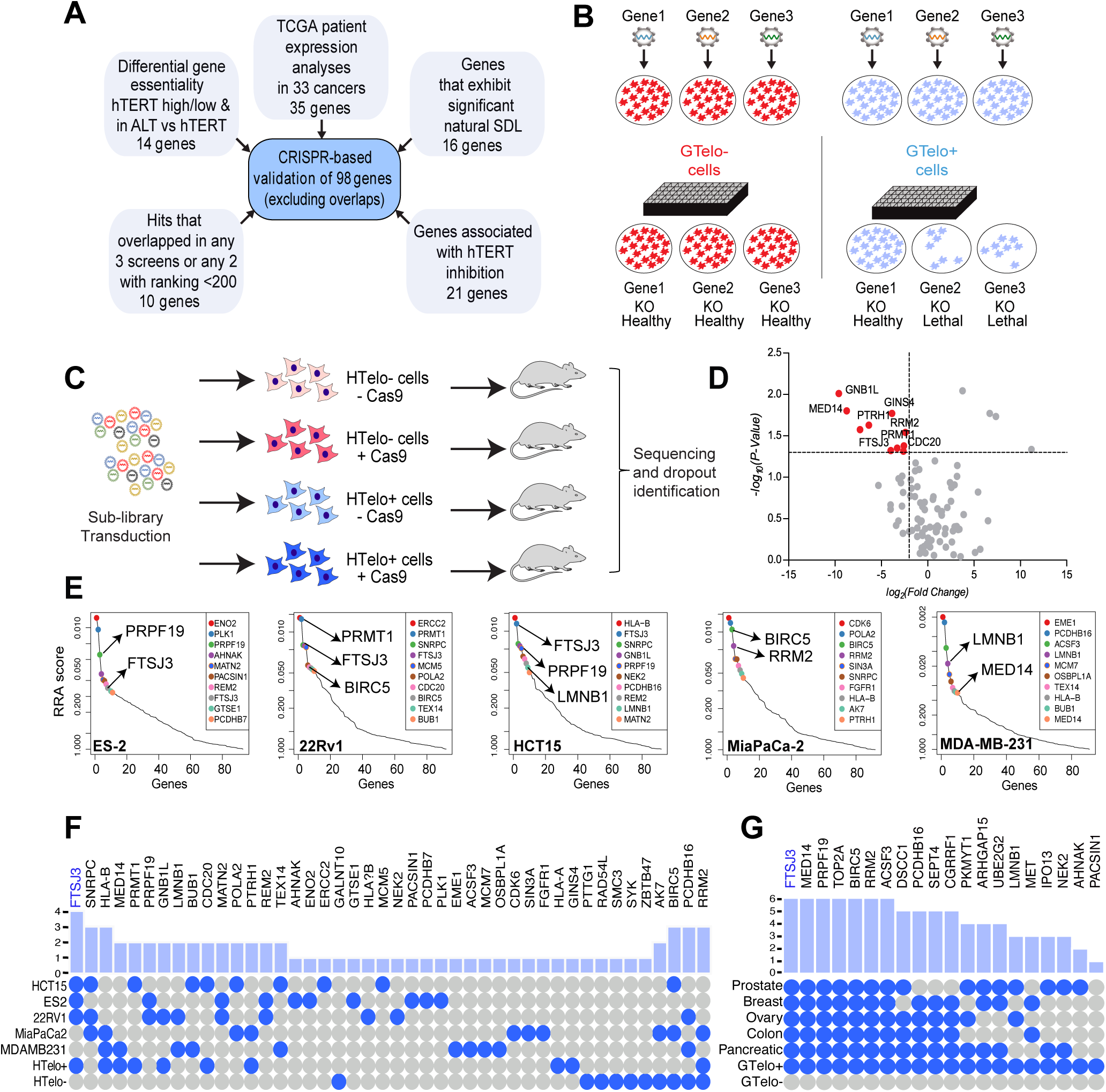
Prioritization and validation of SDL hits using arrayed *in vitro* and pooled *in vivo* CRISPR/Cas9 platforms. **(A)** Schematic representation of five different strategies for choosing 98 SDL candidates from four screens. **(B)** Schematic depicting the validation strategy in which an arrayed CRISPR library was used for one-on-one evaluation of the top SDL hits in GTelo-/GTelo+ isogenic models. Briefly, an equal number of Cas9-expressing isogenic cells were seeded in 96-well plates, followed by transduction with lentiviruses expressing two individual sgRNAs targeting each of the SDL hits or control sgRNA in each transduction. The plates were subsequently imaged for 7 days in an IncuCyte S3® platform, after which the cells were quantified to confirm the true SDL hits. **(C)** Schematic depicting the validation strategy using a pooled CRISPR library for the evaluation of the top SDL hits in an *in vivo* HTelo+/HTelo-isogenic model. **(D)** Volcano plot showing the results of the pooled *in vivo* CRISPR screen comparing HTelo+ cells to HTelo-cells. For statistical analysis, a t-test in conjunction with a permutation test was used to calculate the p values. The names of some of the top ten SDL hits are highlighted. **(E)** MAGeCK plots of all the *in vivo* pooled CRISPR screens were conducted in five xenograft models. The plots highlight the top ten SDL hits in each screen identified by Robust Ranking Aggression scores. **(F)** Summary dot plots of the top SDL hits from *in vivo* pooled CRISPR screen. The blue dots represent the SDL hits that were validated in the specified model. **(G)** Summary dot plots of the top SDL hits from the *in vitro* arrayed CRISPR validations.

For validation, we used an arrayed *in vitro* CRISPR/Cas9 strategy in GTelo-/GTelo+ cells and a pooled *in vivo* CRISPR/Cas9 strategy in HTelo-/HTelo+ cells. Nontargeting sgRNAs served as controls. *In vitro*, 53 of 98 SDL hits preferentially induced lethality in GTelo+ cells (p < 0.05), while 18 showed a similar trend but were below significance **(Fig. S4A-B)**. From this, we selected the top 20 SDL hits for further validation in 10 hTERT^positive^ cell lines representing breast, colon, pancreatic, ovarian, and prostate cancers **(Fig. S4C-D)**. Matching CRISPR knockouts were introduced, cell proliferation was monitored for seven days, and the obtained data confirmed SDL properties of the hits **(Fig. S5A-B)**. Genes well known for their roles in telomere maintenance were among the top hits. For example, BIRC5 (Survivin), an inhibitor of apoptosis, and LMNB1, a nuclear lamina component, have established links to telomere regulation^24,25^, while RRM2 inhibition has already been shown to impact hTERT^positive^ over ALT cells preferentially^26^. Notably, the BIRC5 inhibitor YM155 selectively suppressed hTERT^positive^ cells **(Fig. S2B)**. Collectively, these findings validate our SDL hits and highlight their potential as selective targets in hTERT-driven cancers.

To further validate these hits *in vivo*, we employed a pooled CRISPR/Cas9 screen using a sub-library of 196 sgRNAs targeting the 98 SDL hits in HTelo-/HTelo+ cells. Transduced tumor cells were implanted into mice, and after reaching ∼500 mm^3^, tumors were harvested and sequenced for sgRNA dropout. Genes like FTSJ3, MED14, RRM2, GNB1L and PRMT1 emerged as top SDL hits in hTERT^positive^ tumors **(Fig. 2C-D)**. Expanding this *in vivo* screen to multiple hTERT-overexpressing models (breast, pancreatic, ovarian, colon, and prostate cancers) once again identified FTSJ3, MED14, PRMT1, GNB1L, PRPF19, and LMNB1, although some SDL hits were tissue-specific **(Fig. 2E-F)**. BIRC5 and RRM2 were among the top hits, but their effects on HTelo-cells reduced their prioritization **(Fig. 2F)**. By comparing hits from cell-based assays **(Fig. 2G)** with those from *in vivo* pooled CRISPR screens **(Fig. 2F)**, we prioritized FTSJ3 as the top candidate for further investigation.

### FTSJ3 is essential in hTERT^positive^ tumor-agnostic cancer models

To explore the essentiality of FTSJ3 in tumors with elevated hTERT expression, we rigorously assessed the dependency of hTERT^positive^ cells on FTSJ3 using five independent model systems. Our study encompassed six hTERT^low^ non-malignant cell lines (Hs895.SK skin fibroblasts, PNT1B human prostate epithelial cells, HGF-1 gingival fibroblasts, HPDE pancreatic duct epithelial cells, HFF-1 foreskin fibroblasts and BJ foreskin fibroblasts), four hTERT^low/negative^ ALT cell lines (U-2 OS, KMST1, NCI-H1573, and SK-LU-1), and four hTERT^positive^ cancer cell lines (22Rv1, MDA-MB-231, ES-2, and MIA PaCa-2). Additionally, we included two primary pancreatic cancer cell lines (PaCaDD-115 and PaCaDD-119) and a patient-derived pancreatic cancer organoid model (PANC-004). RT-qPCR confirmed that hTERT expression was several folds higher in hTERT^positive^ models compared to the GTelo-cell line **(Fig. 3A)**, validating our comparative approach. All models also expressed FTSJ3 **(Fig. S6A)**.

**Figure 3:**
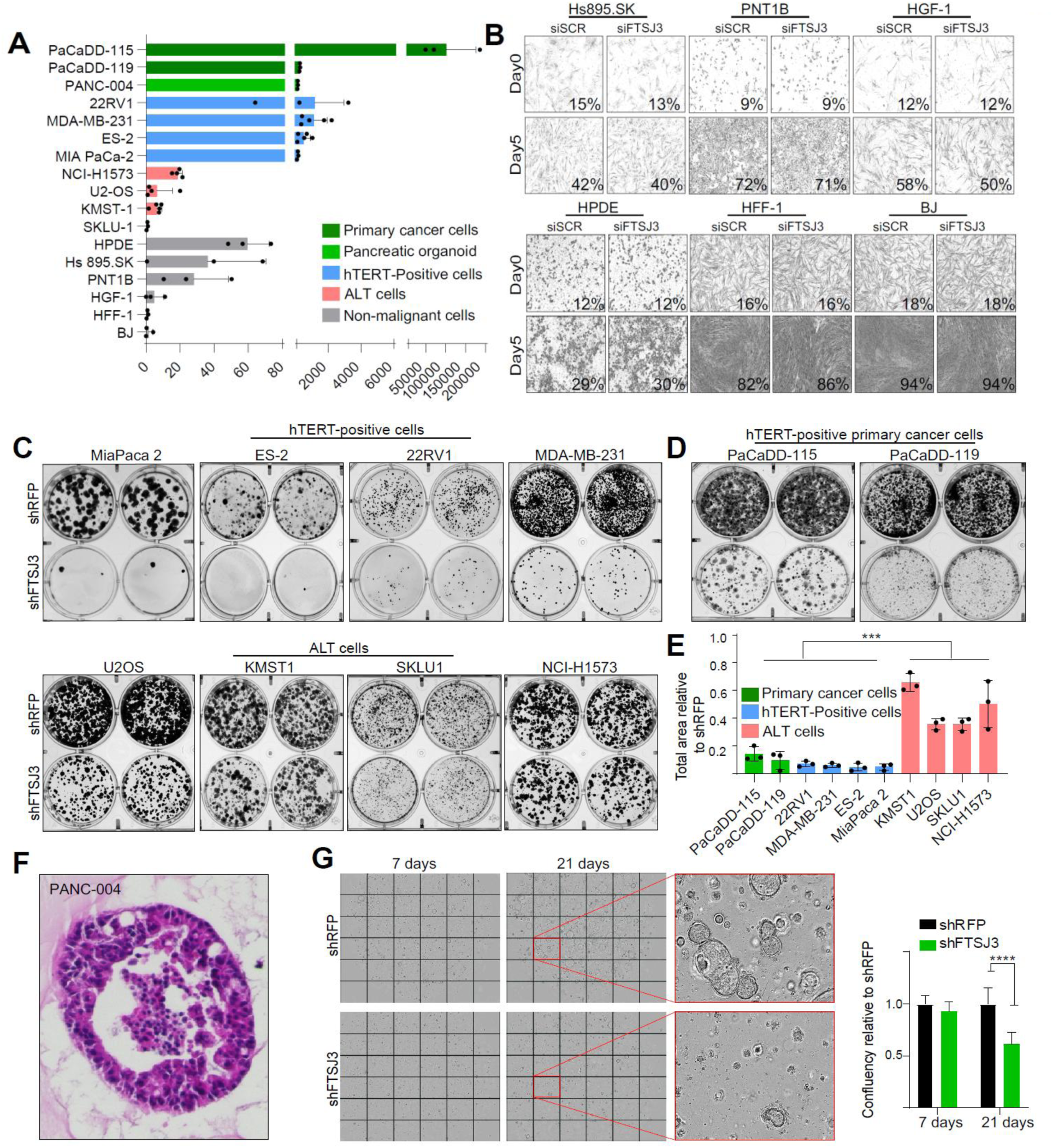
FTSJ3 is preferentially essential in hTERT^high^ tumor-agnostic cancer models. **(A)** Bar graph of the qPCR data showing fold change in hTERT gene expression values relative to GTelo-expression. Primary pancreatic cancer cells and a pancreatic cancer organoid are shown in green, hTERT-positive cells are shown in blue, ALT-positive cells are shown in red, and non-malignant cells are shown in gray. N=3-5 biological replicates. **(B)** Representative images of non-malignant cells captured by the IncuCyte imaging system before transduction with siRNA (day 0) and five days after transduction. The percentages shown in each panel represent the percentage of cell confluency as calculated by the IncuCyte analysis tool. Colony formation analysis was not performed for these cell lines, as they failed to produce colonies. **(C)** Colony formation assay was performed in hTERT-positive (MIA PaCa-2, ES-2, 22Rv1, and MDA-MB-231) and ALT (U-2 OS, KMST1, SK-LU-1, and NCI-H1573) cell lines following FTSJ3 knockdown. Transduction with shRFP was used as a control. The Image J software was used to quantify colonies’ abundance by measuring total areas of the colonies in each well. The data are presented as fold change relative to matching shRFP controls. **(D)** Colony formation assay was performed in hTERT-positive primary cancer cells following FTSJ3 knockdown, as in (C). **(E)** Bar graph quantifying colonies in 3C and 3D. Ordinary one-way ANOVA with Tukey’s multiple comparison test; **p < 0.0024, N=3 biological replicates. **(F)** Hematoxylin and Eosin-stained histology section of original pancreatic tumor (left) and organoid (right). * Represent annotation of perineural invasion. **(G)** PANC-004 organoid images were captured with the IncuCyte 7 days and 21 days after shFTSJ3 transduction. The graph illustrates confluency relative to the shRFP, quantified by the IncuCyte imaging software. RM two-way ANOVA, with Sidak’s multiple comparisons test, ****p < 0.0001, N=6.

FTSJ3 knockdown had no significant negative effects on non-malignant cell lines **(Fig. 3B)**. In contrast, FTSJ3 knockdown markedly reduced the colony formation ability of hTERT^positive^ cancer cell lines and primary pancreatic cancer cell lines, while ALT-positive cell lines were only minimally affected **(Fig. 3C-E)**. This was further corroborated in a patient-derived pancreatic cancer organoid model, which maintained a gland-forming histology like the original tumor **(Fig. 3F-G)**. The effects on the organoid model suggest that FTSJ3 loss may mitigate tumor heterogeneity, reflecting the diverse cellular composition of the patient’s tumor. The efficiency of FTSJ3 knockdowns was confirmed through RT-qPCR and Western blotting **(Fig. S6B-C)**. Overall, our findings highlight the critical role of FTSJ3 in supporting the viability of hTERT^positive^ tumors, suggesting that targeting FTSJ3 may offer a promising therapeutic strategy for overcoming the challenges posed by tumor heterogeneity.

### FTSJ3 deficiency disrupts telomere anchoring via heterochromatin modulation

To elucidate the mechanism underlying the SDL relation between *FTSJ3* and *hTERT*, we investigated the potential role of FTSJ3 in telomere maintenance and telomerase activity. In these experiments, telomere shortening in response to FTSJ3 suppression was observed in hTERT^positive^ MIA PaCa-2 cells at multiple time points, compared to U-2 OS ALT cells (**Fig. 4A).** Universal STELA confirmed an increase in short (≤ 5 kb) and critically short (≤ 2 kb) telomeres in FTSJ3-silenced MIA PaCa-2 cells. In contrast, U-2 OS ALT cells showed no such changes (**Fig. 4B)**. Notably, telomerase activity remained unaffected by FTSJ3 loss **(Fig. 4C)**, suggesting that FTSJ3 influences telomere stability independent of telomerase.

**Figure 4:**
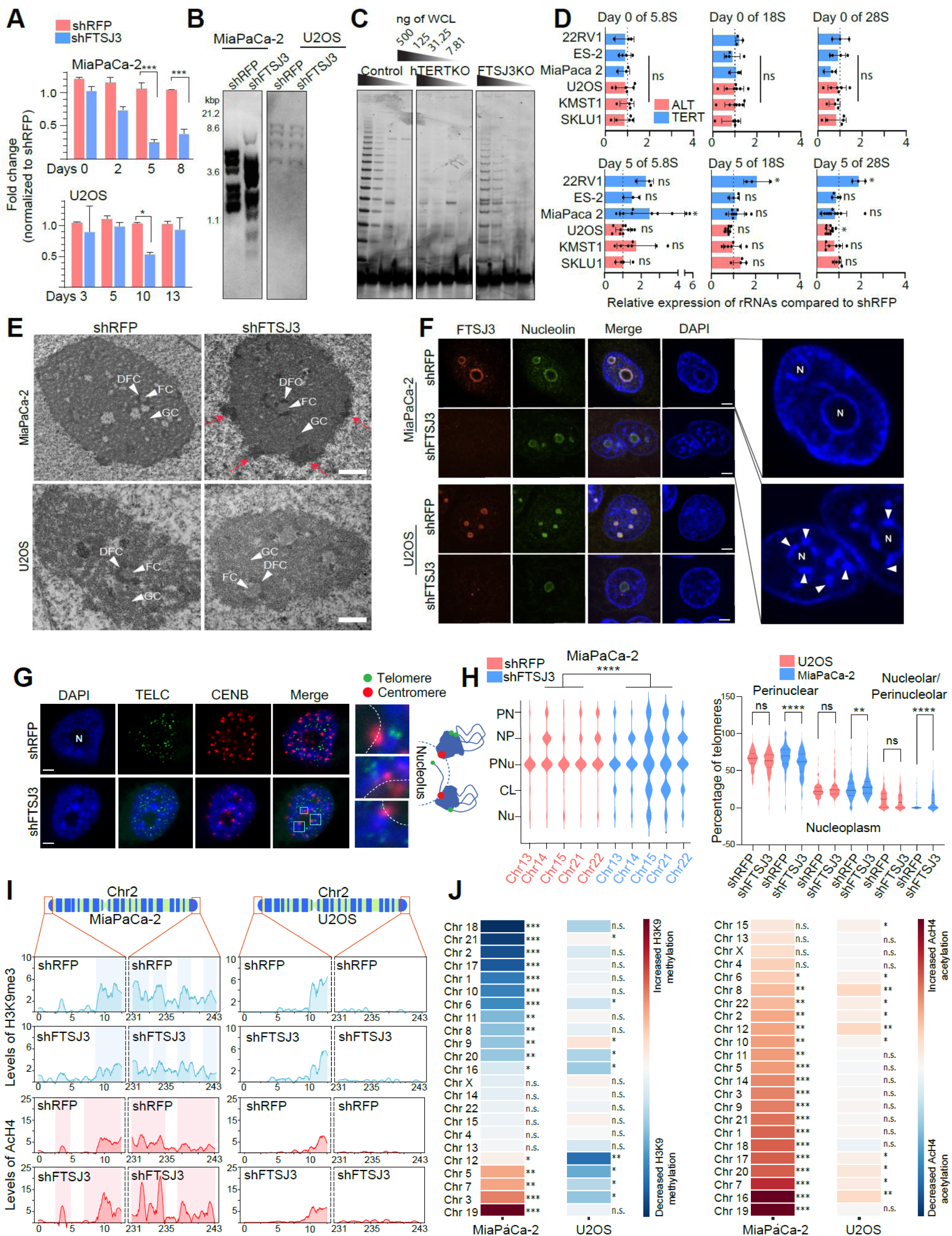
FTSJ3 regulates heterochromatin formation. **(A)** Quantitative PCR was used to measure telomere length in both MIA PaCa-2 cells and U-2 OS cells. Multiple t-tests, N=3, *p < 0.01; ***p < 0.001. **(B)** Representative Universal STELA image showing an increase in short telomeres in shFTSJ3-knockdown MIA PaCa-2, but not in U-2 OS cells. **(C)** TRAP assay measuring telomerase activity in FTSJ3 knockout MIA PaCa-2 cells compared to that in nontargeting control and hTERT knockout cells. **(D)** Bar graph showing relative expression of key rRNA subunits in FTSJ3-silenced cells compared to shRFP treated cells with no consistent significant deviations from the baseline. **(E)** Electron micrographs of MIA PaCa-2 and U-2 OS transduced with shRFP and shFTSJ3, 5 days following puromycin selection. FC: fibrillar center; DFC: dense fibrillar component; GC: granular component. Scale bar: 1µm. **(F)** Representative immunofluorescence images of MIA PaCa-2 and U-2 OS transduced with shRFP or shFTSJ3. FTSJ3: red; nucleolin: green; nucleus: DAPI. N: nucleolus. Arrows indicate heterochromatin aggregates. Scale bar: 5µm **(G)** Representative immune-FISH images of MIA PaCa-2 transduced with shRFP or shFTSJ3. Nucleus: DAPI; telomere TELC: green; centromere CENB: red. Teal squares highlight regions where the centromere PNA probe colocalizes with heterochromatin aggregates but not with the telomere probe. The broken line illustrates the nucleolar periphery. DAPI is overlayed on TELC and CENB to illustrate chromocenters in shFTSJ3. Scale bar: 5µm. **(H)** Quantitation of centromeric positioning (left) and telomeric positioning in MIA PaCa-2 cells transduced with shRFP or shFTSJ3. PN: perinuclear, NP: nucleoplasm, PNu: perinucleolar, CL: chromocenter-like, Nu: nucleolar. The chi-square test for trend was used for the left panel statistics, while a non-parametric Mann-Whitney U test was used for the right panel. N=50-60; ****p < 0.001. **(I)** Subtelomeric H3K9me3 trimethylation (upper, blue) and subtelomeric AcH4 acetylation (lower, red) profiles of MIA PaCa-2 and U-2 OS for chromosome 2. The 5% regions most distal to both chromosome arms are shown. Highlighted regions in the MIA PaCa-2 panels show areas of decreased methylation (upper) or increased acetylation (lower) in shFTSJ3 samples. The x-axes show the base position in Mbp, and the y-axes show the methylation scores. The data represent the mean methylation and acetylation values across two biological replicates (N=2). Peaks were smoothed with a Savitzky–Golay filter using a 100 bp window and a first-degree polynomial. **(J)** Heatmap showing sum of methylation (left) or acetylation (right) scores across the distal 5% of each arm for all chromosomes in the indicated cell type. p-values were calculated using the Mann–Whitney U test between the shRFP and shFTSJ3 samples. ***, p < 0.001; **, p < 0.01; *, p < 0.5; ns, not significant.

Given the FTSJ3’s reported role in ribosome biogenesis through one of three pathways in the maturation of 18S rRNA precursors^27^, we tested whether this function contributed to its selective lethal effects on hTERT^positive^ cells. However, silencing FTSJ3 did not have any consistent selective impact on the expression of 5.8S, 18S, or 28S rRNAs in either hTERT^positive^ or ALT cells **(Fig. 4D)**, indicating that rRNA synthesis is unlikely to explain the observed SDL effects. Since nucleolar integrity is crucial for ribosome biogenesis, alterations in nucleolar structure can indicate disruptions in ribosome production^28^. Therefore, we examined nucleolar integrity using electron microscopy and observed well-defined fibrillar centers (FC), dense fibrillar components (DFC), and granular components (GC) in both hTERT^positive^ and ALT cells, indicating a preserved nucleolar organization post-FTSJ3 knockdown **(Fig. 4E)**. The presence of these distinct, well-organized nucleolar sub-compartments indicated that the nucleolar organization is intact. The spatial separation of the various stages of ribosome biogenesis is preserved, suggesting that FTSJ3 depletion does not disrupt nucleolar structure or ribosome assembly pathways **(Fig. 4E**-white arrows). These results suggest that the ribosome biogenesis function of FTSJ3 is unrelated to its SDL properties in hTERT^positive^ cancer cells.

Interestingly, we found aberrant heterochromatin aggregation at the perinuclear and perinucleolar regions in ∼70-80% FTSJ3-deficient hTERT^positive^ MIA PaCa-2 cells compared to ∼20-30% of matching control and ALT cells (p < 0.001) **(Fig. 4E**-red arrows). DAPI staining confirmed the presence of these aggregates in multiple models, resembling ‘chromocenters’, where satellite DNA within heterochromatin clusters together, typically near the nuclear envelope or nucleoli^29^. **(Fig. 4F**; **Fig. S7A)**. To determine whether these chromocenter-like aggregates (CLA) were linked to centromeres (pericentromeric heterochromatin, PCH) or telomeres (sub-telomeric/telomeric heterochromatin, STH), we performed FISH using centromeric (CENB) and telomeric (TELC) probes. CLA was predominantly associated with centromeres, with telomeric signals on their periphery, suggesting that they may represent collapsed chromosomal arms, lacking proper telomeric anchoring typically occurring at the nuclear peripheries^30,31^ **(Fig. 4G).** Specifically, the centromeres of acrocentric chromosomes localized to these CLA, and the overall telomere distribution showed gross migration towards the nucleolus **(Fig. 4H**; **Fig.S7B-C)**.

Both PCH and STH exhibit a complex chromatin landscape, characterized by high levels of histone H3 Lys9 trimethylation (H3K9me3) and low histone H4 acetylation (AcH4), associated with repetitive genomic regions. We found H3K9me3 epigenetic modifications enriched within CLA, while H4Ac remained peripheral in FTSJ3-deficient hTERT^positive^ cells **(Fig. S8A)**. Chromatin immunoprecipitation followed by sequencing (ChIP-seq) analysis revealed a significant decrease in H3K9me3 within STH regions in FTSJ3-deficient hTERT^positive^ cells (p < 0.05), accompanied by increased AcH4 levels (p < 0.05) **(Fig. 4 I-J; Fig. S8B**). The ALT models did not show such drastic changes **(Fig. 4 I-J; Fig. S8B)**. PCH regions showed no significant changes **(Fig. S9)**. Combined with previous reports that H3K9me3 positions heterochromatin at the nuclear periphery^31^, our results suggest that reduced H3K9me3 disrupts telomere anchoring, leading to telomere migration towards the nucleolus. This indicates that FTSJ3 is critical for maintaining heterochromatin stability and proper telomeric positioning via H3K9me3 in hTERT^positive^ cells. Although chromatin modifications can occur as part of the DNA damage response (DDR), the changes observed in FTSJ3-deficient cells were independent of DNA damage. Specifically, only ATM phosphorylation was increased in hTERT^positive^ cells without other detectable DDR signaling events, which matched previously reported DDR-independent ATM action^32^ **(Fig. S10A-B)**. Together, these results indicate that the loss of FTSJ3 affects the epigenetic status of STH regions, highlighting its critical role in stabilizing heterochromatin and telomere anchoring in hTERT^positive^ cells.

### The FTSJ3/TERRA/SUV39H1 axis is critical for heterochromatinization in hTERT^positive^ cells

A decrease in H3K9me3 levels caused by the loss of FTSJ3 within the STH regions of hTERT^positive^ cells, but not in ALT models, suggests distinct mechanisms for heterochromatin maintenance between these two cell types **(Fig. 4 I-J; Fig. S8B)**. Among the mammalian family of H3K9 histone methyltransferases (HMTs), SUV39H1 emerged as a potential key player, as it not only promotes H3K9me3 but also stabilizes heterochromatin independently of its methyltransferase activity^33^. Chromatin immunoprecipitation followed by qPCR (ChIP-qPCR) revealed a marked reduction in SUV39H1 binding to telomeric ends in FTSJ3-deficient hTERT^positive^ cells **(Fig. 5A)**, correlating with the observed decrease in H3K9me3 levels. This prompted us to consider whether SUV39H1 is predominantly involved in maintaining heterochromatin in hTERT^positive^ cells. Analysis of TCGA gene expression data from patient samples indicated that both FTSJ3 and SUV39H1 are significantly overexpressed in hTERT^positive^ cancers, but not in those patients lacking hTERT expression **(Fig. 5B)**. Moreover, a significant positive correlation (Spearman correlation, r = 0.21, p = 2.3 × 10^-69^) in expression between FTSJ3 and SUV39H1 was observed only in hTERT^positive^ cancers but not in those patients lacking hTERT expression (Spearman correlation, r = 0.053, p = 0.038) **(Fig. 5C).** Given that ALT cells strongly rely on SETDB1, another HMT, for H3K9me3 maintenance^34^, these observations suggest that different HMTs could be employed for heterochromatin stability in hTERT^positive^ versus ALT cells. Therefore, we next focused on understanding how FTSJ3 might influence SUV39H1 recruitment to STH regions to maintain heterochromatin stability in hTERT^positive^ cells.

**Figure 5:**
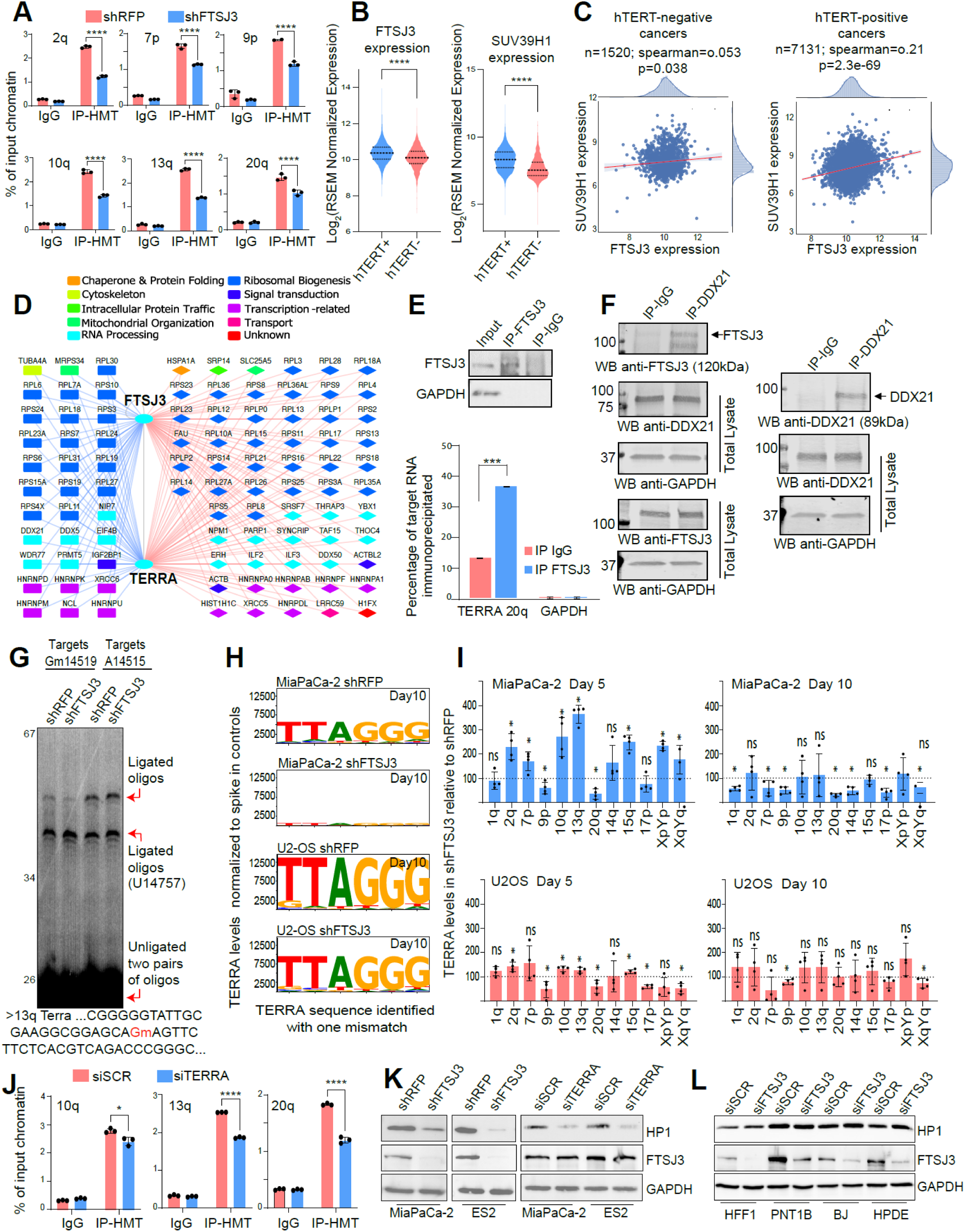
FTSJ3 is required for 2’-O-methylation of TERRA. **(A)** ChIP-qPCR showing the physical interaction between SUV39H1 and subtelomeric region in control and FTSJ3 knockdown cells. Quantification of the immunoprecipitated DNA was normalized to input. Two-way ANOVA, ***p< 0.001. **(B)** The violin plot shows mRNA expression levels of FTSJ3 and SUV39H1 in TERT-positive patients (blue, n = 7,131) and TERT-negative patients (representing ALT) (red, n = 1,520). Statistical significance was determined using the non-parametric Mann-Whitney U test, with p-values less than 1.00×10^-9^ for both comparisons. **(C)** The correlation between FTSJ3 and SUV39H1 in TERT-negative (representing ALT) (n = 1,520) and TERT-positive (n = 7,131) patients was assessed using Spearman rank correlation. Each dot represents a patient sample, and the red line indicates the linear trend. In TERT-negative samples, the correlation coefficient was r = 0.053 with a p-value of 0.038, while in TERT-positive patients, the correlation coefficient was r = 0.21 with a p-value of 2.3 × 10⁻⁶⁹. **(D)** Commonly shared physical interactions between FTSJ3 and TERRA as derived from Simabuco *et al*.^56^, and Scheibe *et al*.^59^. Interactions shown with blue edges (left) are those genes that also exhibit SDL with hTERT. Protein names are color-coded using their broad GO SLIM terms. **(E)** RNA immunoprecipitation (IP) shows the physical interaction between FTSJ3 and TERRA (Input: 20% of the total sample). TERRA levels in FTSJ3 immuno-precipitates and IgG controls were detected via qPCR. Quantification of the percentage of immunoprecipitated TERRA RNA based on total RNA. Two-way ANOVA, ***p < 0.001. **(F)** Western blot assessing FTSJ3 and DDX21 presence in total lysates as well as in samples after immunoprecipitation using anti-DDX21. **(G)** In the first two lanes, the ligation reaction contained two pairs of oligos targeting Gm14519 and a control site, U14757 (unmodified). In the last two lanes, oligo pairs targeting A14515, which is not 2’-O-methylated (negative control), and the control site (U14757) were used. The upper band is the ligation product of a pair of oligos targeting Gm14519 or A14515, and the lower band is the ligation product of the control pair of oligos targeting U14757. **(H)** TERRA transcripts were quantified via Oxford Nanopore Technologies’ MinION platform, allowing one mismatch, demonstrating the number of telomeric repeats normalized against spike-in control. **(I)** TERRA levels were measured on chromosomes 1q, 2q, 7p, 9p, 10q, 13q, 14q, 15q, 17p, 20q, XpYp, and XqYq in both MIA PaCa-2 cells and U-2 OS cells at two different time points (day 5 and day 10) after FTSJ3 knockdown. The dotted lines represent TERRA levels in matching shRFP controls. Note that although ALT cells have higher TERRA levels, these graphs were normalized with their corresponding shRFP controls. Multiple t-tests, *p < 0.01; ns - not significant. N=4 biological replicates. **(J)** ChIP-qPCR shows the physical interaction between SUV39H1 and subtelomeric region in control (siSCR) and TERRA knockdown cells. Quantification of the immunoprecipitated DNA was normalized to input. Two-way ANOVA, *p<0.05, ****p< 0.0001. **(K)** Western blot analysis measuring HP1-alpha levels in MIA PaCa-2 and ES-2 cells five days following shFTSJ3 transduction (left) or five days post siTERRA transfection (right). siSCR: Scrambled control. **(L)** Western blot analysis measuring HP1-alpha levels in non-malignant cell lines five days following siFTSJ3 transfection. siSCR: Scrambled control.

Since non-coding RNAs like TERC and TERRA play crucial roles in telomere synthesis and heterochromatin formation^35–37^, we hypothesized that FTSJ3, being an RNA 2’-O-methyltransferase, might regulate SUV39H1 recruitment by modifying these RNAs. Specifically, we first considered whether FTSJ3 could influence telomeric chromatin indirectly via hTERC methylation, given FTSJ3’s known interaction with nucleolin^38^ and its predicted association with hTERC-binding proteins^39^ **(Fig. S10C)**. This hypothesis was grounded in the idea that modifications to hTERC could potentially alter the structure or stability of the telomeric chromatin complex, thereby influencing SUV39H1’s access to telomeric regions. Although hTERC was found to be 2’-O-methylated at ∼120 nucleotides from the 5’ end, FTSJ3 depletion did not alter hTERC methylation **(Fig. S10D)**, nor did it affect telomerase activity **(Fig. 4C)**. These results indicated that hTERC was not a part of the pathway through which FTSJ3 could be modulating heterochromatin and SUV39H1 recruitment. This outcome led us to shift our focus to TERRA, another essential non-coding RNA directly interacting with telomeric chromatin.

Analysis of FTSJ3 and TERRA interaction networks^38,40^ revealed a significant overlap in their binding partners, with a direct interaction between FTSJ3 and TERRA being the most notable **(Fig. 5D)**. Further reinforcing FTSJ3 as an important hTERT-specific SDL hit, we found several shared binding partners of both TERRA and FTSJ3 as SDL hits in our genome-wide screens, including the RNA helicases DDX21 and DDX5 **(Fig. 5D)**, both of which interact with TERRA^41,42^. RNA immunoprecipitation and co-immunoprecipitation experiments confirmed a direct interaction between FTSJ3 and TERRA **(Fig. 5E)**, as well as FTSJ3’s association with DDX21, but not DDX5 **(Fig. 5F**; **Fig. S11A)**. Supporting this, DepMap gene essentiality scores^23^ of known ALT and hTERT cell lines **(Table S5)** revealed that DDX21, but not DDX5, exhibited an SDL relationship with hTERT **(Fig. S11B)**. In summary, the direct interaction between FTSJ3 and TERRA, along with its association with DDX21 and the resulting impact of FTSJ3 deficiency on SUV39H1 recruitment and H3K9me3 levels, points towards a critical role for FTSJ3/TERRA/SUV39H1 (FTS) axis in STH stability in hTERT^positive^ cells.

### Loss of FTSJ3 affects TERRA methylation and TERRA levels

Based on our findings indicating the role of FTSJ3 – TERRA interaction, we set out to investigate whether FTSJ3 loss affects putative TERRA methylation, particularly concerning heterochromatin regulation. To assess this, we performed a 2’-O-methylation assay^43^, which revealed a ∼60% reduction in methylation at position Gm14519 on TERRA from chromosome 13q following FTSJ3 knockdown **(Fig. S11C)**. A ligation-based assay^43^ confirmed that G14519, but not the control site A14515, was methylated in an FTSJ3-dependent manner **(Fig. 5G)**. In cells with intact FTSJ3, G14519 methylation reached ∼70%, dropping to ∼25% after FTSJ3 silencing. These results conclusively demonstrate that FTSJ3 mediates 2’-O-methylation of TERRA at G14519 in the 13q region. Our results do not preclude the likelihood that FTSJ3 silencing may also impact other 2’-O-methylation sites in TERRA. The presence of short, repeated TERRA sequences in the primer extension assay resulted in multiple bands, posing challenges in detecting additional 2’-O-methylation sites.

To further investigate the functional consequences of FTSJ3-dependent TERRA methylation, we employed nanopore sequencing using the MinION platform **(Fig. S11D)**. After normalizing to spike-in controls, in agreement with their roles in the ALT recombination mechanism, we observed significantly higher TERRA levels in ALT cells than hTERT^positive^ MIA PaCa-2 cells **(Fig. 5H)**. Importantly, FTSJ3 knockdown resulted in a 10-fold reduction in TERRA levels in hTERT^positive^ cells, while ALT models exhibited only a 2-fold decrease **(Fig. 5H)**. Interestingly, our direct measurement of TERRA abundance by chromosome end-specific RT-qPCR demonstrated an initial increase in TERRA levels in hTERT^positive^ cells, which was consistent with previously reported TERRA upregulation in response to telomere shortening^44^. Remarkably, at day 10, this was followed by a significant reduction in TERRA levels, specifically in hTERT^positive^ cells. In contrast, only modest reductions were observed in U-2 OS cells, consistent with increased basal TERRA levels in ALT models^34^ **(Fig. 5I)**. These findings establish that FTSJ3 controls TERRA methylation and its abundance in hTERT^positive^ cells.

### FTSJ3 and TERRA safeguard STH in cancer cells as their loss leads to telomeric defects and genome instability

Our results indicate that FTSJ3 directly interacts with TERRA to mediate its 2′-O-methylation and regulate its abundance. We performed TERRA knockdown experiments to determine whether the defective heterochromatin phenotype caused by decreased SUV39H1 association with telomeres in FTSJ3-deficient cells is TERRA-dependent. Similar to FTSJ3 loss, TERRA depletion led to a partial reduction in SUV39H1 binding at telomeric regions, suggesting TERRA’s role as a key mediator in the FTS axis **(Fig. 5J)**. These findings strongly support a model in which FTSJ3 maintains telomeric heterochromatin stability through a TERRA-dependent mechanism, ensuring proper SUV39H1 recruitment and sustaining H3K9me3 levels in hTERT^positive^ cells. Supporting this model, previous studies have shown that TERRA interacts with SUV39H1 at damaged telomeres^45^ and that TERRA depletion is associated with reduced H3K9me3 and impaired recruitment of heterochromatin protein 1 (HP1) to telomeres^36^. Since H3K9me3 is a binding platform for HP1^46^, we hypothesized that FTSJ3 modulates H3K9me3 levels within the STH by regulating TERRA, thereby influencing SUV39H1 recruitment. Consistent with this hypothesis, we observed reduced HP1 protein levels in both FTSJ3-deficient and TERRA-silenced hTERT-positive cells **(Fig. 5K)**. These results suggest that FTSJ3’s regulation of TERRA is critical for SUV39H1 loading at telomeric ends, thereby maintaining stable STH in hTERT^positive^ cells.

Given that *de novo* telomere repeat synthesis and subsequent heterochromatinization occur exclusively in hTERT^positive^ cells, we next asked whether HP1 levels are affected by FTSJ3 loss in non-malignant, hTERT^negative^ cells. Using four non-malignant cell lines, we found that HP1 levels remained unchanged upon FTSJ3 knockdown **(Fig. 5L)**. This indicates that FTSJ3’s role in regulating HP1 and telomeric heterochromatin stability is selective for hTERT^positive^ cells, and that normal cells are not affected by FTSJ3 suppression, as they are not actively engaged in the synthesis of telomere repeats, further highlighting the therapeutic potential of our work.

TERRA has also been shown to recruit members of nucleolar remodelling complex (NoRC)^47^ and polycomb repression complex (PRC2)^36^ to telomeres to establish STH. We next asked if this function of TERRA is also dependent on FTSJ3. Indeed, we observed that SMARCA5, a component of NoRC, localized to CLAs in FTSJ3-deficient cells, though it did not accumulate at peripheries of the CLA **(Fig. S12A)**, while PRC2 recruitment to telomeres remained largely unaffected by FTSJ3 silencing **(Fig. S12B)**, indicating that the disruption of TERRA regulation in FTSJ3-deficient cells impairs specifically the recruitment of select heterochromatin maintenance factors, such as SMARCA5 and SUV39H1, but does not broadly impact other chromatin-modifying complexes like PRC2 **(Fig. S12B)**. These findings suggest that FTSJ3-mediated modulation of TERRA is crucial for recruiting heterochromatin-stabilizing factors to stabilize STH regions, particularly in the chromosomes of hTERT^positive^ cells.

Given that defective heterochromatin integrity correlates with genomic instability^48^, we conducted telomere-FISH combined with the TeloView imaging platform^49^. Consistent with our mechanistic findings, FTSJ3 silencing significantly reduced total telomere intensity, indicating telomere shortening **(Fig. 6A, B**; **Fig. S12C)**. This effect was evident in MIA PaCa-2 cells but not ALT cells **(Fig. S12D)**, implicating FTSJ3 loss in genome instability, specifically in hTERT^positive^ cells. R-loops are representative of the overall genome instability phenotype^50^ and indeed, dot blots of DNA:RNA hybrids demonstrated FTSJ3 deficiency in MIA PaCa-2 cells led to a strongly increased accumulation of R-loops **(Fig. 6C),** which was not necessarily restricted to telomeric regions. In contrast, R-loop levels in ALT cells remained unchanged with FTSJ3 silencing, highlighting a critical distinction in heterochromatin regulation between hTERT^positive^ and ALT cells **(Fig. S12E).** Specifically, ALT-positive cells maintain high basal R-loop levels that can serve as initiation signals for recombination-dependent telomere maintenance through the ALT mechanism^51^.

**Figure 6:**
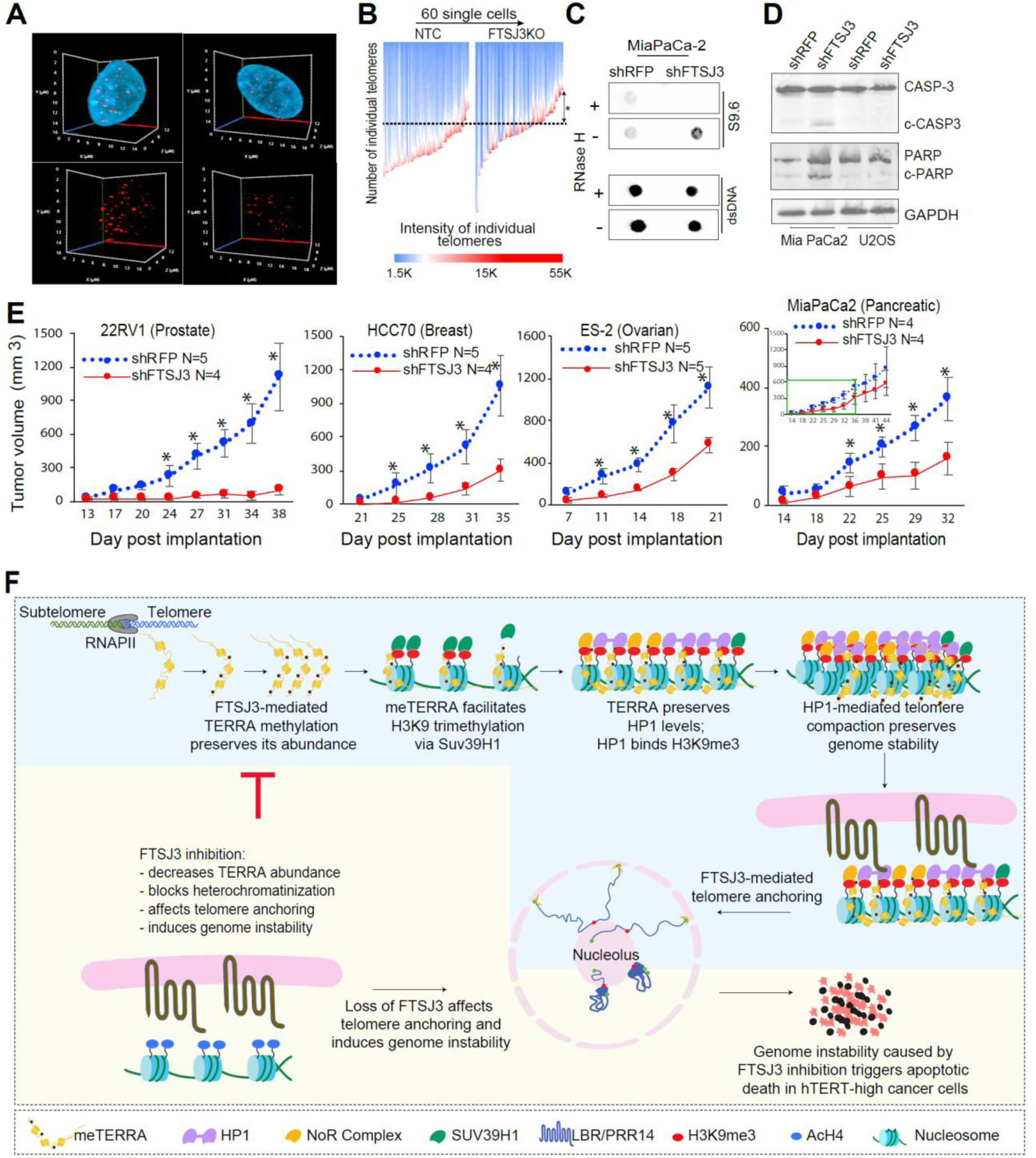
Loss of FTSJ3 induces genome instability and reduces tumor growth. **(A)** Representative images of 3D telomere organization in the NTC and FTSJ3 knockout MIA PaCa-2 cells with 3D nuclei (blue) and 3D telomeres (red) are shown in the top panel. The bottom panel represents the spatial representation of the telomeres. **(B)** Gradient scale representation of the number of telomeres and their corresponding telomere staining intensities gathered in 60 random NTC and FTSJ3 knockout MIA PaCa-2 cells. The Y-axis represents the number of telomeres measured in each cell, and the color code represents the intensity values. **(C)** FTSJ3 was silenced in MIA PaCa-2 cells, and dot blot analysis was subsequently performed to quantify the DNA:RNA hybrids. Equal amounts of DNA were spotted on nitrocellulose membranes, and R-loops were detected using the S9.6 antibody. RNaseH treatment was also conducted, and the resulting product was used as a negative control to confirm the specificity of the signal. The ssDNA signal was used as an internal sample loading control. **(D)** Western blot demonstrating that MIA PaCa-2 cells five days after treatment with shFTSJ3 are undergoing apoptosis through Caspase and PARP cleavage. CASP-3: caspase; c-CASP-3: cleaved caspase; c-PARP: cleaved PARP. **(E)** Validation of the tumor-suppressing effects of shRNA-mediated silencing of FTSJ3 (shFTSJ3) in hTERT^high^ mouse xenograft models generated with the pancreatic cancer cell line MIA PaCa-2, prostate cancer cell line 22Rv1, breast cancer cell line HCC70 and ovarian cancer cell line ES-2 using RFP-targeting shRNA (shRFP) as a non-targeting control. Unpaired heteroscedastic t-test, *p < 0.05, N=4-5 mice. **(F)** Proposed mechanism of FTSJ3-loss-mediated toxicity in TERT-positive cells. FTSJ3 methylates TERRA transcripts to stabilize *de-novo* synthesized telomere repeats through heterochromatin formation. FTSJ3 loss prevents this modification, which has downstream implications of decreasing overall TERRA abundance, promoting epigenetic modifications which limit heterochromatinization of the telomeres, and finally resulting in telomere erosion and subsequent genome instability.

Genomic instability is closely linked to the induction of apoptosis^52^ and consistent with this RNA-seq analysis of FTSJ3-deficient hTERT^positive^ cells revealed the upregulation of genes involved in apoptosis. In contrast, ALT cells did not display such significant alterations **(Fig. S13)**. Consistent with this, apoptotic events, including cleaved Caspase 3 and PARP, were significantly upregulated exclusively in hTERT^positive^ cells following FTSJ3 depletion **(Fig. 6D)**, suggesting that FTSJ3 silencing was affecting these cells by inducing apoptotic cell death. To assess the biological relevance of our findings, we examined the effect of FTSJ3 silencing in xenograft models of several human malignancies. Excitingly, these experiments revealed that suppression of FTSJ3 activity consistently reduces tumor growth in hTERT^positive^ prostate, breast, and ovarian cancer models, though the effect was partial in the pancreatic cancer xenograft **(Fig. 6E**; **Fig. S13C)**, highlighting its potential as a therapeutic target in multiple cancer types. Consistent with this, analyses of TCGA data showed better survival of hTERT^positive^ patients with lower FTSJ3 **(Fig. S13D)**

## Discussion

Direct inhibition of telomerase in cancer cells has been pursued through various strategies, with disappointingly limited clinical success to date^53^. This study focused on SDL interactions linked to telomerase expression rather than directly targeting telomerase, identifying FTSJ3 as a critical vulnerability. By leveraging SDL interactions, we expect to overcome the limitations of traditional anti-telomerase strategies. Importantly, unlike conventional target discovery, which often relies heavily on combinations of genomic screenings with literature-based knowledge, we adopted a systematic, unbiased approach in our work. Through an integrated combination of *in vivo* pooled CRISPR screens and *in vitro* validation across diverse models, we rigorously prioritized FTSJ3 as a highly promising therapeutic target. As demonstrated in our validation work, this approach ensures that our findings are robust and broadly applicable across cancer types, positioning FTSJ3 as a central SDL partner in telomerase-associated pathways.

Mechanistically, our findings demonstrate that FTSJ3 methylates TERRA, regulating its abundance and stabilizing STH in hTERT^positive^ cells. Loss of FTSJ3 leads to a relaxed, less stable telomeric heterochromatin structure, which likely impairs telomere anchoring and causes chromatin disorganization, including the formation of perinucleolar CLAs. This structural instability is likely due to the inefficient loading of SUV39H1 at telomeric ends, as FTSJ3-mediated regulation of TERRA is critical for proper SUV39H1 recruitment. This agrees well with the findings of the Lingner lab, which show that the N-terminus of SUV39H1 binds directly to TERRA repeats^45^ and has direct TERRA interaction with telomeric DNA^54^. Therefore, when the loss of FTSJ3 leads to a reduction in TERRA levels, SUV39H1 fails to accumulate at STH, reducing H3K9me3 levels and impeding the recruitment of HP1-alpha, which functions to compact and stabilize heterochromatin. This disruption further destabilizes the heterochromatin structure, promoting genomic instability and enhancing the risk of chromosomal aberrations **(Fig. 6F)**.

Inner nuclear membrane proteins, such as lamin B receptor (LBR), and nuclear lamina-associated proline-rich protein PRR14 are known to anchor heterochromatin to the lamina through HP1 interactions^30,55^. Therefore, the reduction of H3K9me3 and the failure of HP1 recruitment suggest that the attachment of telomeric heterochromatin to the nuclear membrane may be compromised in FTSJ3-deficient cells, which we also observed in our experiments. This chromatin disorganization and CLA formation in FTSJ3-deficient cells further destabilize telomere integrity and triggering apoptotic responses **(Fig. 6F)**. These findings underscore the critical role of telomeric heterochromatinization for maintaining telomere integrity, specifically in hTERT^positive^ cells, where continuous telomerase-driven telomere elongation requires simultaneous establishment of a tightly regulated heterochromatic state. The interaction between FTSJ3 and TERRA likely accounts for the selective suppression of hTERT^positive^ cells following FTSJ3 loss. In contrast, normal cells, which do not actively synthesize telomeric repeats, remain unaffected by the lack of FTSJ3 activity.

Our study also reveals a stark contrast between responses of hTERT^positive^ and ALT cells to FTSJ3 inhibition. ALT cells resist FTSJ3 absence due to their reliance on a recombination-based mechanism for telomere maintenance^7^, which operates independently of FTSJ3-mediated TERRA regulation. The permissive chromatin state in ALT cells facilitates telomeric recombination, bypassing the stringent heterochromatin requirements seen in hTERT^positive^ cells. Moreover, ALT cells utilize a different HMT, SETDB1, to maintain their H3K9me3 status^34^, while our results indicate that hTERT^positive^ cells strongly rely on SUV39H1 for this purpose. This highlights the crucial role of the FTSJ3/TERRA/SUV39H1 (FTS) axis in heterochromatin maintenance in hTERT^positive^ cells, distinguishing them from ALT cells. This differential regulation further emphasizes the specificity of FTSJ3 as a target for hTERT^positive^ cancers while sparing ALT cells and minimizing off-target effects on normal cells.

From a therapeutic standpoint, FTSJ3 represents an attractive target, as its methyltransferase activity, mediated by a conserved S-adenosylmethionine (SAM)-binding domain, is amenable to pharmacological inhibition^56^. This domain, essential for the methylation reaction, is an attractive candidate for small-molecule inhibitors. Additionally, recent studies show that FTSJ3 inhibition can elicit antitumor immune responses^57^. Our findings support this notion, as FTSJ3 affects the loading of SUV39H1 on telomeric DNA, whose inhibition has been linked to enhanced antitumor immunity^58^. However, SUV39H1 is associated with DNA methyltransferases and may influence methylation patterns across multiple genomic regions^59^. Its broad regulatory control raises concerns that its inhibition could lead to unintended effects on gene expression and cellular function, complicating therapeutic applications. In contrast, targeting FTSJ3 could facilitate dual-action therapies: disrupting telomere maintenance in cancer cells to promote their elimination while enhancing antitumor immunity. This positions FTSJ3 as a unique target capable of destabilizing telomeres and amplifying the immune response, offering a compelling strategy for combination therapies in cancer treatment.

Given its pivotal role in telomere maintenance and the potential for pharmacological inhibition, the rapid onset of genomic instability following FTSJ3 loss is particularly significant, especially in the context of telomerase reactivation, a hallmark of most cancers^60^. While telomerase reactivation is often considered a means to restore genomic stability, allowing cancer cells to thrive, the acute genomic instability triggered by FTSJ3 inhibition may exceed this stabilizing influence, effectively preventing cancer progression.

In conclusion, our findings reveal FTSJ3 as a potent and highly specific therapeutic target in hTERT^positive^ cancers. By disrupting TERRA’s methylation, FTSJ3 inhibition drives STH destabilization, leading to selective genomic instability in telomerase-dependent cancer cells **(Fig. 6F)**. As a druggable enzyme essential for telomerase-driven cancers, FTSJ3 offers broader and more feasible translational potential compared to the challenges of directly targeting TERRA. These insights open new avenues for precision therapies that exploit a critical vulnerability in hTERT^positive^ cancers, offering the potential to halt tumor progression at a key moment of instability.

## Methods

### Resource availability

#### Lead contact

Further information and requests for reagents and resources should be directed to and will be fulfilled by the lead contact, Franco Vizeacoumar.

#### Materials availability

This study did not generate new unique reagents.

#### Data Availability

The deep sequencing data from CRISPR screens (.fastq.gz files), raw microarray data (.CEL files) from the shRNA screens, *in vivo* pooled CRISPR screens, Nanopore sequencing and ChIP-seq data have been uploaded to the Gene Expression Omnibus (GEO) database, and the corresponding accession number is <u>GSE211313</u>. All other data reported in this paper will be shared by the lead contact upon request. This paper does not report original code.

### Experimental model and study participant details

#### Cell lines and cell culture

All the cell lines used in this study are listed in **Table S6.** All the cell lines were incubated at 37°C in 5% CO2. The cell lines were either authenticated by cell line authentication services at Genetica DNA Laboratories (Cincinnati, Ohio, USA) or purchased from commercial suppliers. Purchases were made through Cedarlane labs (Burlington, Ontario, Canada), a Canadian distributor for the American Type Culture Collection or from MilliporeSigma. The cell lines purchased from ATCC/Sigma were passaged for less than three months at a time following resuscitation; therefore, no additional authentication was performed. Mycoplasma tests were routinely conducted.

#### Mouse models and tumor xenograft studies

The mice were housed 5 per cage at 23-25°C in a humidity-controlled colony room, maintained on a 12 h light/dark cycle (08:00 to 20:00 light on), with standard food and water provided *ad libitum* and environmental enrichments. All animals were handled in accordance with the approved protocols by the University of Saskatchewan Animal Research Ethics Board (AREB). All mice used in this study were NSG (NOD.Cg-Prkdscidll2rg) and littermates within the same cage were randomly allocated to experimental or control groups.

*In vivo* pooled CRISPR screens were executed as previously described^61^. For the *in vivo* pooled CRISPR screen, tumors were generated by injecting 4 x 10^6^ MIA PaCa-2 cells, 2 x 10^6^ ES-2 cells, 1 x 10^6^ 22Rv1 cells, 3 x 10^6^ HCT-15 cells, 2 x 10^6^ MDA-MB-231 cells or 5 x 10^6^ HT1080 cells, while xenograft tumors were established by injecting 4 x 10^6^ MIA PaCa-2 cells, 1 x 10^6^ ES-2 cells, 1.5 x 10^6^ HCC70 cells, 1 x 10^6^ 22Rv1 cells in 50 μL of PBS, and 50 μL of Matrigel (Corning) into the flank regions or mammary fat pads of ∼6–8-week-old male (MIA PaCa-2, 22Rv1, HCT-15, HT1080, HCC70) and female (ES-2, MDA-MB-231) animals, respectively. Each experimental group consisted of 5 mice, totaling 60 mice for the *in vivo* CRISPR screen and 40 mice for the xenograft tumors. Tumor measurement was initiated when the tumors became palpable. Measurements were performed with a digital caliper twice weekly for at least 20 days. The experiments were terminated, and the animals were sacrificed according to the AREB regulations.

### Methods details

#### Lentivirus production and generation of Cas9 stable cell lines

Lentiviral particles expressing shRNA, sgRNA or Cas9 were generated by transfecting logarithmically dividing HEK-293T cells with the packaging plasmids psPAX2 and pMD2. G with pLKO.1^62^ or Lenti-PB^63^. Transfection was performed in 10 cm plates with media containing 540 μL of Opti-MEM (Gibco, Life Technologies) and 36 μL of X-tremeGENE DNA Transfection Reagent (Roche). The media was changed after 18 hr and replaced with DMEM containing 20% w/v bovine serum albumin (Sigma), after which the viral particles were collected after 24 and 48 hr. Viral harvests were pooled and centrifuged at 1000 rpm for 3 min and stored at -80° C for subsequent use. Stable Cas9-blast cell lines were generated by transducing each cell line with the lentiviral Cas9-blast at a final concentration of 8 μg/mL polybrene for 24 hr. Blasticidin-containing media was changed for every 2 to 3 days until the untransduced cells showed complete cell death.

#### Pooled shRNA and CRISPR/Cas9 screening and data deconvolution

Screening was performed as previously described^15,64^. Briefly, cells were transduced at a low MOI (< 0.3) to avoid multiple infections and maintain at least 200-fold hairpin/guide representation. Cells were collected every 3 days for a minimum of 10 generations, and genomic DNA was extracted using a QIAamp Blood Maxi Kit (Qiagen) at a minimum of three time points. Microarray analyses were performed for shRNA screens, and deep sequencing was performed for CRISPR screens. Primers used for the shRNA library and CRISPR library are listed in **Table S7.** Overall, three time points for each screen with replicates were analyzed by taking the weighted change in the cumulative difference between each time point. In the case of shRNA screening, the signal intensity from the microarray was normalized via quantile normalization, and shRNAs with signals below the background (i.e., log2 scale of less than 8) at the initial time point T0 were removed prior to further analyses to compute the fitness score. For CRISPR/Cas9 screens, FASTQ sequences were aligned to the library sequences using the count function of the MAGeCK software package^65^. Prior to aligning the fastq files, the FASTQC package performed a quality check. The count data were then preprocessed using the ‘CRISPRcleanR’ package to eliminate gene-independent responses to CRISPR‒Cas9 targets, as previously described^14^. Each replicate at each time point for every screen was independently parsed through the CRISPRcleanR package before the corrected counts were used to identify SDL hits. The corrected read counts were then cyclic loess-normalized prior to further analyses to compute the fitness score. The shRNA-or sgRNA-weighted differential cumulative change (*WDC_h/g_*) between hTERT^high^ and hTERT^low^ cells was calculated for each consecutive time point using the following formula:

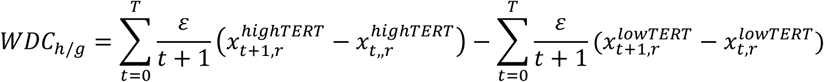

where *x^lowTERT^_t,r_* is the normalized signal intensity of the hTERT^low^ cells at time point *t* ∈ (0, . . *T*) in replicates *r* ∈ (1. . *N*). Similarly, *x^highTERT^_t,r_* represents hTERT^high^ cells. ε is a constant that determines the weight between each time point so that shRNA or sgRNA drops at earlier time points are ranked before the shRNA or sgRNA drops at later time points. Gene level WDC_gene_ was computed as the average of the top two shRNAs/sgRNAs with the most negative values for that gene using the formula below.

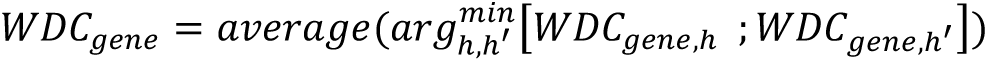

To identify shRNAs or sgRNAs and their corresponding genes that are significantly different between hTERT^high^ and hTERT^low^ cells, Student’s t-test was used in combination with the permutation test p value by estimating the frequency of randomized, shuffled WDCs with more negative values in comparison with the observed gene-level WDC value, as previously described^15^. Bayesian analysis of the gene essentiality algorithm was used to evaluate the performance of the screens^66^. Pooled screening using *in vivo* models was conducted, and the data were analyzed as previously described^61^.

#### Screening data quality analysis

To ensure the performance of the shRNA and CRISPR/Cas9 screens, we used the Bayesian Analysis of Gene Essentiality (BAGEL) software package^67^. The precision–recall curves were calculated based on the core essential and nonessential genes. The F-measure was calculated and found to be > 0.7. For the CRISPR/Cas9 screens, the count summary file from MAGeCK revealed that the Gini coefficients were acceptable, with all the screens having a rating < 0.11 and a mapping read percentage between 55% and 62%^68^. For each screening group, the experiment showed consistent Pearson correlations calculated at the intensity level for the shRNA screens and at the count level for the CRISPR/Cas9 screens ranging between 0.85 and 0.99, suggesting that the replicates were highly reproducible.

#### Computational analyses of TCGA data, DepMAP and CancerRXgenes

To calculate the clinical significance of the gene in relation to telomerase activity, we mined the TCGA-identified normalized gene expression data and analyzed patients with high TERT expression (greater than the median expression of TERT in normal samples for the same tissue type) as well as low SDL hit gene expression (lower than the median expression of the SDL hit in the normal samples of the same tissue type); the remaining patients composed the second group. Then, we calculated the log-rank test to determine whether the patients who exhibited a natural SDL pattern were significantly different (p < 0.05) from the non-SDL-exhibiting group of patients. For SUV39H1 or FTSJ3 expression analyses in patient samples, patients with an RNASeq Expectation Maximization value of 0 for TERT expression were classified as TERT-negative patients, representing ALT patients. Out of the 24 types of tissues studied, two types (READ and THYM) had no ALT patients and were removed from the study. This process left us with data from 22 tissue types, including 1,520 TERT-negative patients (representing 17.6% of all patients) and 7,131 TERT-positive patients (representing 82.4% of all patients). This organized dataset was used to create graphs showing differences and relationships in gene activity for further study. For survival analysis, both TERT-negative and TERT-positive patients were split into two groups based on their FTSJ3 gene activity: high and low. The split was made using the median value of FTSJ3 activity across all patients. A Kaplan-Meier survival analysis was then performed to see how FTSJ3 activity affects patient survival. The statistical significance of differences between these groups was assessed using the Wilcoxon rank-sum test. A p-value of <0.05 was considered statistically significant. For pathway analyses, SDL hits from the shRNA and CRISPR screens were combined for each model system (GTelo and HTelo) separately, and used Gene Set Enrichment Analysis (GSEA, RRID:SCR_003199) was used^69^. We used version 7.4 of the Reactome Molecular Signature Database (MSigDB v.7.4) for GSEA. To further identify the pathways enriched in all four screens, we used the iDep online web platform to determine the pathway enrichment in the Reactome dataset^70^. We used the essentiality score from DepMap for gene essentiality analyses, where gene essentiality was measured for all the genes across multiple cancer cell lines. Using the Cancer Cell Line Encyclopedia dataset, which contains gene expression data for all the genes in each cancer cell line, we classified cell lines into high and low quartiles according to TERT gene expression. Then, for each hit that was identified in our shRNA/CRISPR screens, we calculated a nonparametric Mann–Whitney U test between the essentiality scores of the two classifications. Genes with a significant (p < 0.01) difference between these groups were prioritized for validation studies. The public domain CancerRXgenes database has a compendium of cancer drugs tested across multiple cancer cell lines. We used these data to group cell lines with low or high (top 1%) TERT expression and measured the half-maximal inhibitory concentration (IC_50_) of multiple drugs across these groups of cell lines. For each drug in their dataset, we performed a nonparametric Mann–Whitney U analysis of the difference in the IC_50_ between the 2 groups.

#### Arrayed CRISPR/Cas9 secondary screen and genome editing assay

Selected hits from the screen were validated in an arrayed format. A minimum of two sgRNAs were used in a pooled format to increase editing efficiency. Briefly, glycerol stocks of bacterial cultures harboring each sgRNA of interest were inoculated into deep 96-well blocks containing 1.3 mL of LB media supplemented with 100 µg/mL ampicillin. The plates were sealed with permeable membranes and incubated at 37°C and 300 rpm overnight. The plates were centrifuged at 1800 × g for 10 min, and the plasmids were isolated using a GenElute HP 96-well Plasmid Miniprep Kit (Sigma-Aldrich). Lentiviruses were generated using HEK293T cells (ATCC) in a 96-well format as described above. For hit validation in each cell line, cells were seeded in 96-well plates, and each well received lentivirus from two pooled sgRNAs targeting a single gene. Puromycin selection at 2 µg/mL was conducted for 48 hr to eliminate uninfected cells. Next, the medium was replaced with fresh medium, and the plates were transferred to an IncuCyte S3® Live-Cell Analysis System (Essen BioScience). The cells were left to grow for 7 days to compare the difference in growth between the knockouts and the control, and the media was changed every 48 hours. Genome editing was assessed using a GeneArt® Genomic Cleavage Detection Kit (Thermo Fisher). Briefly, the chromosomal loci of interest were amplified via PCR. Primers were designed with Tm > 55°C. at a length of 18–22 bp with a 45–60% GC content. The primers used in the study are listed in **Table S7**. Primers were designed to amplify sequences approximately 600 bp upstream and 300 bp downstream of the sgRNA target site. The cell lysates (2 μL) were mixed with 1 μL of 10 μM forward or reverse primer mixture (final concentration of 200 nM) and 25 μL of AmpliTaq Gold® 360 Master Mix, and the final volume was increased to 50 μL with water, as recommended by the manufacturer’s protocol. Amplification PCR with denaturing and reannealing reactions was performed at 95°C for 5 min, 95°C–85°C–2°C/sec, 85°C–25°C, and 0.1°C/sec, after which the samples were cooled to 4°C. The heteroduplex DNA generated was digested by adding 1 μL of detection enzyme (endonuclease) followed by incubation for 1 hr at 37°C. The final product was separated on a 2% agarose gel.

#### TeloView analyses to evaluate abnormalities associated with telomeres

The TeloView experiment was conducted as previously described^71^. Briefly, MIA PaCa-2-Cas9 knockout cells were seeded on sterile slides and placed in an incubator at 37°C overnight. The medium was removed, and the slides were washed, fixed, and treated with Triton X-100 and 20% glycerol. After the freeze–thaw process, the slides were washed again with PBS and incubated in fresh 0.1 M HCl. The slides were equilibrated with formamide, and a PNA FISH Kit/Cy3 Probes (Dako) hybridized with telomeres was applied to the selected area. The slides were covered with a coverslip and sealed with rubber cement. The nuclei were counterstained with 4′,6-diamidino-2-phenylindole (DAPI), and the coverslips were mounted using VECTASHIELD (Vector Laboratories). The slides were kept at 4°C until imaging. A total of 60 cells were imaged for nuclei per condition using an AxioImager Z2 microscope (Zeiss). A 63×/1.4 oil objective lens was used for image acquisition. Sixty z-stacks were acquired at sampling distances of x, y: 102 nm and z: 200 nm for each stack slice. ZEN Microscopy Software (Zeiss) was used for 3D image acquisition and processing (the constrained iterative algorithm was used for deconvolution). Deconvolved images were analyzed using the TeloView v1.03 software program^49^ (Telo Genomics). TeloView loads 3D images and determines telomeric signal intensity (telomere length), the number of telomeric signals and telomere aggregates. The constrained iterative algorithm option was used, and at least 30 nuclei were analyzed for each time point.

#### Universal STELA analyses

The universal STELA was performed as described by Bendix *et al*.^72^. Briefly, 1 µg of DNA was digested by a 1:1 mixture of the restriction enzymes MseI (New England Biolabs) and NdeI (New England Biolabs) at 37°C for 1 h followed by 20 min of inactivation at 65°C. Ten nanograms of digested DNA was mixed with 50 μmol 42-mer and 50 μmol 11 + 2-mer oligos **(Table S7)**. The mixture was spun down to decrease the temperature from 65°C to 16°C for 1 h. Twenty units of T4 DNA ligase (New England Biolabs) were added with T4 DNA ligase reaction buffer and left overnight at 16°C. After the overnight incubation, 20 U of T4 DNA ligase and 10^−3^μm Telorette were added, and the reaction mixture was supplemented with T4 DNA ligase reaction buffer and left again overnight at 35°C. The next day, the mixture was inactivated for 20 min at 65°C. Telomere-specific PCR was carried out in a 12 μL volume containing 20–50 pg of ligated DNA, 1× Failsafe PCR PreMix H (Biosearch Technologies), 0.1μm teltail and adapter primers, and 1.25 U of Failsafe enzyme (Biosearch Technologies). The reaction was carried out on a thermocycler under the following conditions: 1 cycle of 68°C for 5 min; 1 cycle of 95°C for 2 min; 26 cycles of 95°C for 15 s, 58°C for 30 s and 72°C for 12 min; and 1 cycle of 72°C for 15 min. Telomere repeat fragment detection was carried out with the Southern technique described for TRF using a (TTAGGG)n DIG-labeled probe and the TeloTAGGG™ Telomere Length Assay Kit from Millipore Sigma.

#### TRAP assay for measuring telomerase activity

The PCR-based telomerase rapid amplification protocol (TRAP) measured telomerase activity. Whole-cell lysates (WCLs) from KO or transfected cells were extracted, and the protein content was normalized via the Bradford colorimetric assay. TRAP was performed using 500, 125, 31.25, and 7.81 ng (4-fold dilution series) of WCL. Telomerase 6-nt telomeric repeats were added at 30°C for 1 hr in the presence of 100 ng of M2 primer, 1× TPCR buffer (20 mM Tris-HCl pH 8.0, 1.5 mM MgCl_2_, 68 mM KCl, 0.05% Tween 20, and 1 mM EGTA), and 2.5 nmol of dNTPs. The reactions were heat-inactivated at 95°C for 3 minutes. The primer extension products were then PCR amplified using 2.5 units of Pfu DNA polymerase in 1x TPCR buffer with 100 ng of the Cy5 fluorophore-labeled reverse primer (CX3), 90 ng of the NS primer, and 5.0 × 10 3 copies of the NSCX3 internal control oligo. The products were resolved on a 10% PAGE gel (10% acrylamide/bis (19:1), 0.5x TBE, 0.1% APS and 0.1% TEMED) and visualized using a Typhoon imager with a Cy5 670BP30 filter (wavelength 655-685 nm).

#### Electron microscopy sample preparation and imaging

Two biologic replicates of MIA PaCa -2 and U-2 OS transduced with shRFP or shFTSJ3 were cultured under puromycin selection for 7 days before cultured cells had growth media aspirated and washed in PBS. Cells were fixed in 2% glutaraldehyde in 100mM NaAsO_2_(CH_3_)_2_ buffered to a pH of 7.3 at 4 °C for 3 h. Fixative was collected and adherent cells were scraped and combined with the collected fixative before being centrifuged at 200G for 5 min at 4 °C. Supernatant was discarded and the cell pellet was subsequently resuspended in 100mM NaAsO_2_(CH_3_)_2_ and incubated at 20 °C for 15 min. Centrifugation and resuspension was repeated before storing the samples at 4 °C for future processing. Samples were again pelleted and then resuspended with warm 1% low melting point agarose before being repelleted and stored. Samples were postfixed in freshly prepared 1% OsO_4_ in 100mM NaAsO_2_(CH_3_)_2_ for 1 h at 20 °C. Samples were then rinsed in 100mM NaAsO_2_(CH_3_)_2_ three times before beginning a serial dehydration and en bloc staining with UO₂(CH₃CO₂)₂ in the following process: 50% ethanol for 5-10 min, saturated UO₂(CH₃CO₂)₂ for 1 h, 70% ethanol for 5-10 minutes, 90% ethanol for 5-10 min, 95% ethanol for 5-10 min, thrice 100% ethanol for 5-10 min, and thrice with propylene oxide for 5 min. Samples were then infiltrated with 1 part Epon/Araldite : 2 parts propylene oxide for 30 min, 2 parts Epon/Alrdite : 1 part propylene oxide for 2 h, and then in pure Epon/Araldite overnight. Samples were then oriented in molds in fresh Epon/Araldite and the resin polymerized at 60 °C for 24-48 h. Ultrathin sections were cut to 90 nm with a diamond knife and placed on 200 Cu grids for imaging. Samples were imaged on a Hitachi HT7700 transmission electron microscope between 2000 and 30,000x magnification.

#### RNA immunoprecipitation (RNA IP)

Cell pellets were obtained and resuspended in RNA IP lysis buffer (980 µL of Pierce^TM^ RIPA buffer (Thermo Scientific), 10 µL of Halt^TM^ protease inhibitor, Thermo Scientific, and 10 µL of RNase inhibitor; Promega). The suspension was incubated on ice for 30 min with intermittent vortexing. Centrifugation at 8000 rpm and 4 ℃ for 30 min was then performed to remove the debris. A total of 1 mg of the extraction was taken for each IP, and 100 µg was taken as the 10% input. Protein A/G beads (Thermo Scientific) were prepared by washing three times with 1 mL of PBS. After each wash, the beads were collected on magnetic plates. 70 μL of protein A/G beads were needed for each IP, and after the final wash, each protein A/G bead prep was resuspended in 100 µL of PBS. A total of 1 mg of the cell lysate, 10 µL of Halt^TM^ protease inhibitor, 10 µL of RNase inhibitor, 2 µg of antibody (IgG negative control and target protein antibody), 100 µL of washed protein A/G beads, and a suitable volume of PBS were added to one tube to reach a total volume of 1 mL for one IP. The mixture was incubated at 4°C overnight on a rotator. The beads were washed three times with PBS the next day, and the final collected beads were resuspended in 100 µL of PBS. A total of 30 µL of the beads was collected for western blotting, and the remaining 70 µL of the beads was used for RNA extraction and qPCR.

#### R-loop dot blot assay

Genomic DNA was extracted from cell pellets using a DNeasy Blood & Tissue Kit (Qiagen), and the extracted DNA (50 µg each) was fragmented using 300 µL of restriction enzyme cocktail containing 1 µL of HindIII (20 U/µL), 1 µL of EcoRI (20 U/µL), 2 µL of BsrGI (10 U/µL), 1 µL of XbaI (20 U/µL), and 4 µL of SspI (5 U/µL) in NEB Buffer 2 (NEB) at 37°C overnight. The mixture was divided into two parts: one was treated with 2 U of RNase III (NEB) and 4 U of RNase T1 (Thermo Scientific) for 2 h, and the other was treated with 4 U of RNase H (NEB), 2 U of RNase III (NEB) and 1000 U of RNase T1 at 37°C for 2 h. The fragmented DNA samples were purified using a DNeasy Blood & Tissue Kit. Each sample was diluted to 250 ng in 50 µL of TE, spotted on a nylon membrane and crosslinked by UV treatment. The membrane was blocked with 5% sucrose and incubated with an anti-DNA‒RNA hybrid antibody, clone S9.6 (Sigma-Aldrich) or dsDNA antibody (as an internal control; Neobiotechnologies) overnight at 4°C, followed by washing three times with TBST and incubation with secondary antibody for 1 h at room temperature. After washing three times with TBST again, the membrane was treated with enhanced chemiluminescence (ECL) reagent (Bio-Rad) and visualized using a ChemiDoc MP Imaging System (Bio-Rad).

#### Primer extension-based 2′-O-methylation assay

To detect the 2′-O methylated nucleotides within TERRA and hTERC, a standard primer extension-based modification assay (with high and low dNTP concentrations) was carried out essentially as previously described^43^. Briefly, ∼7.5 μg of total RNA was mixed with 5′-radiolabeled DNA oligonucleotides (primer), 13q TERRA R or hTERC (151-170) in a volume of 8 μL of annealing buffer (50 mM Tris-HCl pH 8.3, 60 mM NaCl2, 10 mM DTT). The mixture was denatured at 95°C for 2 min, reannealed at 55°C for 3 min, and added to 12 μL of primer extension mixture consisting of 9.72 mM Tris-HCl (pH 8.3), 11.34 mM NaCl, 1.94 mM DTT, 5.94 mM Mg(OAc)2, 1 unit of SuperScript III reverse transcriptase (Thermo Scientific), and 0.4 mM dNTPs (high) or 0.01 mM dNTPs (low). Primer extension was carried out at 42°C for 30 min. The extension products were extracted with PCA (phenol:chloroform:isoamyl alcohol 25:24:1; Thermo Scientific), precipitated with ethanol, resolved on an 8% polyacrylamide-8 M urea gel, and visualized by autoradiography.

#### Ligation-based assay for detecting 2’-O-methylated sites

To identify the exact site of the 2’-O-Me modification in TERRA RNA, we performed a ligation-based assay^43^. First, TERRA RNA was purified from human total RNA using biotinylated DNA oligo (Biotin-TERRA-specificRT (Integrated DNA Technologies)) and a streptavidin–biotin affinity pull-down assay. Briefly, ∼50 μg of human total RNA was mixed with 10 μL of 20 μM biotinylated oligo and 20 μL of 5× NET-2-MgCl_2_ buffer (250 mM Tris-HCl, 750 mM NaCl, 0.25% (v/v) NP-40, 10 mM MgCl_2_, pH 7.5). The total volume was brought to 100 μL with ddH_2_O. The mixture was heated at 95°C for 2 min and gradually cooled at room temperature for 10 min. In the meantime, 100 μL of streptavidin magnetic beads (New England Biolabs) was transferred to a new 1.5-mL Eppendorf tube, and the beads were magnetically collected. After the supernatant was discarded and the beads were washed twice with 200 μL of 1× NET-2-MgCl_2_ buffer (50 mM Tris-HCl, 150 mM NaCl, 0.05% (v/v) NP-40, 2 mM MgCl_2_, pH 7.5), the RNA/biotinylated oligo mixture was added to the beads. The sample was incubated for 1.5 h at room temperature. After washing the beads with 200 μL of 1× NET-2-MgCl_2_ buffer four times, the pulled-down RNA was eluted with 150 μL of dissociation buffer (10 mM Tris-HCl, 0.1% (w/v) SDS, and 0.5 mM EDTA, pH 7.5) at 95°C for 2 min. The supernatant was immediately collected. The elution process was repeated two more times. PCA and ethanol precipitation were carried out to recover the eluted RNA. After washing with 70% ethanol, the RNA pellet was resuspended in 20 μL of ddH_2_O.

To check G14519 and A14515 (with the non-2’-O-methylated site as a control) of 13q subtelomere RNA (chr13: 114339022 - 114354019), the DNA oligo pairs below were used; Gm14519-F with Gm14519-A and A14515-F with A14515-A, where the “A” oligos were 5’ [^32^P]-radiolabeled. In all ligation reactions, a control pair of DNA oligos (control, which binds to an unmodified region ∼200 nts downstream of Gm14519) was also used: CF and CA. The ligation-based assay was performed as described by Huang *et al*.^43^. Briefly, the purified TERRA RNAs and oligo pairs were mixed in 0.2X NET-2-MgCl_2_ annealing buffer (2 μL of TERRA RNA, 1 μL of radiolabeled “A” oligos, and 0.5 μL of 10 μM “F” oligos), brought to 8 μL with ddH_2_O) and heated at 95°C for 2 min. The heated samples were gradually cooled at room temperature for 10 min. Each sample was further supplied with 1 μL of 10X DNA ligase buffer (New England Biolabs) and 1 μL of (1 U/μL) T4 DNA ligase (New England Biolabs). The ligation reactions were carried out at 37°C for 30 min. After ligation, PCA extraction and ethanol precipitation were carried out, and the resulting pellet was resuspended in 5 μL of ddH_2_O. 5 μL of 2× loading dye was subsequently added. The samples were resolved on an 8% denaturing polyacrylamide gel (containing 7 M urea), and the radioactive bands were detected via autoradiography.

#### TERRA-Seq Nanopore Sequencing and Analysis

MIA PaCa-2 and U-2 OS cells were transduced with shRFP and shFTSJ3 virus separately. Day 5 and Day 10 cell pellets were collected and proceeded with RNA extraction. For cDNA synthesis, 2 µg of total RNA was reverse transcribed with TERRA-specific primers using SuperScript III Reverse Transcriptase (Thermo Fisher Scientific) and Template Switching RT Enzyme Mix (NEB). TERRA-seq libraries were constructed using Native Barcoding Kit 24 V14 (Oxford Nanopore Technologies). Sequencing was conducted by MinION Mk1B platform (Oxford Nanopore Technologies).

Nanopore sequencing data were processed using a method adapted from Rodrigues et al.^73^ to investigate sub-telomeric regions across the human genome. Briefly, the process began with the extraction of de-barcoded nanopore reads from the raw sequencing output, which were then assigned to their respective samples after removing barcode sequences. A sub-telomeric reference was created from both ends of each chromosome, using the T2T CHM13v2.0/hs1 genome assembly. This reference was specifically designed to focus on the repetitive regions near the telomeres.

The nanopore reads were aligned to this reference using MiniMap2^74^, a tool optimized for long-read sequencing. The alignment was performed with the parameters ‘-k15 -w5’ (adjusting the minimizers for the query) and ‘-ax map-ont’ (optimized for Oxford Nanopore reads). These settings improved the accuracy of mapping in repetitive regions. After alignment, sequences mapped to the sub-telomeric regions were extracted from the Sequence Alignment Map (SAM) files. The telomeric repeat motif “TTAGGG” was counted, allowing for up to one mismatch to account for potential sequencing errors. This analysis provided insight into the length and composition of telomeric sequences across the genome. To ensure data accuracy, the telomeric repeat counts were normalized against the DNA Control Strand (DCS), a 3.6 kb amplicon targeting the 3’ end of the Lambda genome. The DCS served as a positive control, enabling quality control by accounting for sequencing and alignment biases. The final graphs were generated using the Python logomaker library.

#### RNA extraction, quantitative PCR for TERRA expression, and rRNA quantification

RNA was extracted from the samples with a PureLink RNA Kit (Thermo Fisher). A reverse transcription assay was performed by using the Applied Biosystems™ High-Capacity cDNA Reverse Transcription Kit following the manufacturer’s instructions. Real-time PCR was performed by using PerfeCTa SYBR® Green FastMix PCR Reagent. GAPDH expression was used for normalization, and gene expression was quantified using the 2-ΔΔCt method. A previously established qPCR method was used to measure relative TERRA expression levels^75^. Briefly, total RNA was extracted from ∼3 million cells using a RNeasy Mini Kit (Qiagen) and treated thrice (twice on-columns and once in-solution) with RNAse-free DNase to eliminate genomic DNA contamination according to the manufacturer’s instructions. RNA purity and concentrations were estimated by a NanoDrop® ND-1000 Spectrophotometer (Thermo Fisher Scientific, Waltham, MA, USA). For cDNA synthesis, 3 µg of total RNA was reverse transcribed with TERRA-and GAPDH RT-specific primers using SuperScript III Reverse Transcriptase (Thermo Fisher Scientific). The 1q, 2q, 7p, 9p, 10q, 13q, 14q, 15q, 17p, 20q, XpYp, and XqYq forward and reverse primers were used^75,76^. A 2× Power SYBR Green PCR Master mix (Thermo Fisher Scientific) was used to amplify the samples in triplicate using a StepOne Real-Time PCR System (Applied Biosystems). The data were normalized against GAPDH as a reference gene, and relative changes in expression levels were calculated using the 2^−ΔΔCt^ method. RT‒ PCR was performed using an Applied Biosystem’s High-Capacity cDNA Reverse Transcription Kit for rRNA quantification, with a total of 1500 ng of RNA per 20 µL reaction. cDNA samples were diluted at a 1/5 ratio. Diluted cDNA (3 µL) was combined with 10 µL of Luna Universal qPCR Master Mix (New England Biolabs), and a final concentration of 0.25 µM was used for forward and reverse primers, for a final volume of 20 µL. Real-time PCR primers targeting 18S, 28S, and 5.8 rRNA transcripts were used as described previously^77^. The thermal cycling conditions were as follows: initial denaturation for 60 sec at 95°C; 40 cycles of 15 sec at 95°C and 30 sec at 60°C; and a melting curve. All the reactions were performed in triplicate. The Applied Biosystems StepOnePlus (Thermo Fisher) was used for real-time PCR experiments. The relative expression levels of the rRNAs were evaluated using log2 (2ΔCt) calculations and subsequently relativized to shRFP controls for all the cell lines. GAPDH was used as the reference gene for all the cell lines.

#### Chromatin immunoprecipitation followed by sequencing (ChIP-seq)

The cells were fixed directly on plates with 1% formaldehyde at room temperature for 10 min. Then, 125 mM glycine was added to the media to quench the reaction. After washing with ice-cold PBS twice, the cells were scraped and collected in the EP tube. The cell pellets were resuspended in ChIP lysis buffer (700 µL per 10 M cells, ChemCruz) and sonicated to distribute the sizes of the DNA fragments from 200 bp to 1000 bp. The sonication conditions used differed between the cell types and sonicators, and the chromatin shearing efficiency was checked through reverse cross-linking and AGE. 100 µL lysate was taken for each IP, and 50 µL was taken as the input. For each IP sample, 3 µL of Halt^TM^ protease inhibitor (Thermo Scientific), 3 µL of phosphatase inhibitor (Thermo Scientific), 2∼5 µg of anti-trimethyl-histone H3 (Lys9) antibody (Sigma) or anti-acetyl-histone H4 antibody (Sigma) was added to the lysate, as was a certain volume of RIPA buffer (Thermo Scientific), for a final volume of 300 µL. The IP mixtures were incubated at 4°C overnight on a rotator. 50 µL per sample Protein A/G beads (Thermo Scientific) were added to the mixture after washing three times with 1 mL of PBS the next day, followed by a 1-2 h incubation at 4°C with rotation. The beads were washed with low-salt wash buffer, high-salt wash buffer, and LiCl buffer (BioWorld) three times each. Then, 250 µL of elution buffer (ChemCruz) was added to the beads, which were subsequently incubated at room temperature with rotation for 30 min. The supernatants were combined and transferred to new EP tubes. In each IP tube (500 µL), 20 µL of 5 M NaCl and 4 µL of RNase A (Thermo Scientific) were added, and in each input sample (50 µL of lysate + 70 µL of elution buffer, 120 µL in total), 4.8 µL of 5 M NaCl and 1 µL of RNase A (Thermo Scientific) were added, followed by overnight incubation at 65°C. 4 µL Proteinase K (NEB) per IP mixture and 1 µL per input mixture were added the next day and incubated at 65°C for 1 h. After incubation, the DNA was purified using phenol:chloroform extraction (Invitrogen) and eluted with 20 µL of nuclease-free water. The IPed DNA and input DNA samples were sent to the Next-Generation Sequencing Facility of The Hospital for Sick Children for library construction and sequencing. The ChIP-seq libraries were sequenced through a Novaseq SP Flowcell PE2×100 bp (Illumina).

The sequencing quality was assessed using FastQC version 0.11.9, and no quality issues were found. The adaptors and low-quality bases in the sequencing data were trimmed using atria version 3.2.1 with the default settings. Bowtie2 version 2.5.0 aligned the reads to the human reference genome with the “very sensitive settings” profile. SAMtools version 1.16.1 was used to convert the files into SAM files in BAM files and to sort and index the BAM files. Using the default settings, peaks were identified using epic2, a high-performance reimplementation of SICER. The output bed files were reduced to four columns to create bedgraph files, which were then converted to bigWig files using bedGraphToBigWig. The BigWig files were processed in python using pyBigWig, which is part of the DeepTools software ecosystem. To measure histone modification levels, the most distal 5% on both arms of each chromosome and the 5% on both arms of each chromosome abutting the canonical boundaries of the centromeres were taken for analysis of the subtelomeric and pericentromeric regions, respectively. The data represent the mean peak intensity from two replicates for both methylation and acetylation. The data were assessed for normality using the Shapiro–Wilk test. As the data were not normally distributed, nonparametric statistical tests were used (the Mann–Whitney U test was used for unpaired data, and the Wilcoxon signed-rank test was used for paired data). All the statistical tests were conducted using the python package SciPy. Visualizations were created in python using the packages seaborn, pandas, and matplotlib. When continuous peak data were visualized, a Savitzky–Golay filter using a 100 bp window and a first-degree polynomial was used to smooth the data visualization.

#### Colony formation assay

The cells (2 x 10^3^ cells/well for U-2 OS, 5 x 10^3^cells/well for MIA PaCa-2, ES-2, MDA-MB-231, 1 x 10^4^ cells/well for 22Rv1, KMST1, SK-LU-1, and NCI-H1573) were seeded in 6-well plates and incubated at 37°C under 5% CO_2_ for several days (10 days for MIA PaCa-2, ES-2, MDA-MB-231, 16 days for 22Rv1, U-2 OS, KMST1, SK-LU-1, and NCI-H1573), after which the media was replaced every 3 days. Six-well plates were washed with PBS, fixed with 4% formaldehyde, and stained with 0.5% crystal violet. Colonies were quantified with ImageJ software, and images were taken with the ChemiDoc MP Imaging System (Bio-Rad). For N-methyl mesoporphyrin IX (NMM) treatment, the following modifications were made: MIA PaCa-2, U-2 OS, KMST1, and 22Rv1 cells were seeded at 500, 1000, 1000, and 5000 cells per well, respectively, and cultured for 7–14 days. They were all treated with 2 μM NMM or DMSO every 3 days.

### Quantification and statistical analysis

GraphPad Prism software was used for all tests. A one-tailed unpaired Student’s t-test was used for analysis. The data are presented as the mean ± standard deviation (SD). The significance of our results was determined by setting p < 0.05, and the error was reported as the sum or subtraction of the standard deviation (SD). The nonparametric Mann–Whitney U test was used to compare two groups. Survival analyses were performed using the Kaplan–Meier estimator with the nonparametric log-rank test to measure the equity of strata. To calculate the significance of the difference between the GTelo+ and GTelo-strains in validating the 194 selected genes, we normalized the growth percentage of each SDL partner to that of the nontargeting control (NTC), and Student’s t-test was used to determine the significance of the differences between the groups. The standard error was calculated to compare the means of the replicates between the GTelo+ and GTelo-cells. An unpaired heteroscedastic t-test was used for xenograft work to determine statistical significance. Chi-square analysis was used for TeloView data.

## Acknowledgment

We thank members of the Vizeacoumar and Freywald laboratories for their comments and Mary Kinloch for comments on tumor pathology. Primary prostate epithelial PNT1B cells were a gift from Dr. Michael Cox, University of British Columbia. Primary pancreatic cells are a gift from Drs. Prama Pallavi and Felix Rückert, University Medical Center Mannheim, Heidelberg University, Mannheim, Germany. **Financial Support:** This work is supported by operating grants from the Canadian Institutes of Health Research (PJT-178137; PJT-156401), operating funds from the OvCAN-Cancer Research Society (#2021-OG-876486), CoMRAD funding from the College of Medicine, University of Saskatchewan, and the Be Like Bruce Organization to F.J.V and A.F.; Cancer Research Society (#2014-OG 19190, #2021-OG-837261); the Canadian Foundation for Innovation (CFI-33364); and Saskatchewan Cancer Agency operating grant with funds donated to the Cancer Foundation of Saskatchewan to F.J.V. J.M.Y.W. supported the project from the Cancer Research Society grant funds (#2018-OG-23052). S.M. supported the project through the Canada Research Chairs funds. G.D. supported the work with funds from the Canadian Institutes of Health Research (PJT-156017). F.S.V. is supported by the College of Medicine, U of S. C.G.L. and L.K. were supported by the Mitacs Global Fellowship. A SHRF Fellowship supported R.D.. H.E. and H.P. were supported by the CoMGRAD award, College of Medicine, U of S. J.D.W.P. is supported by a Canada Graduate Scholarship from the Canadian Institutes of Health Research, a Health Sciences Graduate Scholarship from the College of Medicine U of S, and a Doctoral Research Award from the Cancer Research Society (1160056). V.M. is supported by the Canada Graduate Scholarship - Doctoral from the Canadian Institutes of Health Research (FBD-187665) as well as the Health Sciences Graduate Scholarship from the College of Medicine at the University of Saskatchewan.

## Author contributions

Conceptualization: F.S.V., YT.Y., J.M.Y.W., A.F., and F.J.V.; Investigation: J.D.W.P., F.S.V., O.A., Y.Z., V.M., H.A., L.K., A.RP., H.D., L.H.G., P.W., A.G., C.D., T.F., R.D., H.E., A. S., J.P.V., H.P., K.R., K.N., D.M.O., M.LW., A.M.M., A.A., J.L.X., N.A., E.P.M., P.G., J.S., D.D., C.GL., and P.T.; Resources: M.L., G.D., N.J., G.G., A.K., S.A., C.E., K.H.B., Y.W., R.A.DP., S.M., YT.Y., J.M.Y.W., A.F., and F.J.V.; Writing - Review & Editing: all authors; Funding acquisition: G.D., S.M., YT.Y., J.M.Y.W., A.F., and F.J.V.

## Supplementary Figures

**Figure S1: (A)** Left panel: Immunofluorescence of ALT-associated promyelocytic leukemia (PML) bodies in GTelo-/GTelo+ isogenic cell lines. Scale 10 mm. Right panel: Quantitation of the formation of ALT-associated PML bodies in GTelo- and GTelo+ isogenic cells. **(B)** Western blot showing stable Cas9 expression in the GTelo-/GTelo+ and HTelo-/HTelo+ isogenic cell lines. **(C)** Quality assessment of the barcoded products via a bioanalyzer. The quality of the amplified CRISPR amplicons was verified using bioanalyzer gel electrophoresis. None of the samples showed any secondary peak contamination. **(D)** Correlations between replicates in all screens are represented as scatter plots. Pearson correlations were computed between replicates to measure reproducibility, and the corresponding values were provided. **(E)** Venn diagram showing the overlap between the shRNA and CRISPR/Cas9 screens in the GTelo-/GTelo+ and HTelo-/HTelo+ models. As expected from multiple earlier studies, we observed poor overlap between shRNA and CRISPR/Cas9 screens.

**Figure S2: (A)** Cytoscape representation of the SDL hits from four screens arranged based on cellular compartments. The four screens are placed in the four corners, with the SDL genes identified in each of them represented as nodes connected in blue edges. The nodes are color-coded based on Gene Ontology slim terms. The red edges are interactions derived from ingenuity pathway analyses. A large number of nuclear and mitochondrial genes were found to have extensive interactions, as depicted by the red lines derived from the analyses. SDL hits with inhibitors are represented as triangles. **(B)** SDL hits with existing chemical inhibitors were evaluated for preferential suppression of hTERT^high^ cell lines using the CancerRXgene drug database. Cell lines were classified as hTERT^high^ (red) or hTERT^low^ (blue) based on their hTERT expression levels. The sensitivity of the cell lines to the selected inhibitors was plotted as a dose-response curve.

**Figure S3: (A)** Correlation cluster plot for the expression of hTERT with the expression of each screening hit across 33 cancer types. Red indicates a highly positive correlation between hTERT and the SDL gene. The magnified list of genes represents SDL hits that are co-overexpressed with hTERT in multiple cancer types. Gene names are provided on the side, and each column represents a cancer type in the TCGA, and the abbreviations follow TCGA nomenclature. **(B)** Gene essentiality scores derived from DepMap data were used to validate SDL hits. Cell lines were classified based on hTERT expression levels (left) or as ALT and TERT lines (right), and gene essentiality scores were plotted to determine whether SDL hits are preferentially essential for hTERT^high^ cell lines. The top SDL hits that showed significant differences in essentiality between the groups are presented. Nonparametric Wilcoxon rank sum test, *p < 0.01; ***p < 0.0001. **(C)** Left panel: schematic representation of the method for prioritization of clinically relevant SDL hits. Right panel: Kaplan–Meier plots of SDL hits in four different cancer types from TCGA are presented. Bottom panel: A few examples of specific SDL candidates associated with better patients’ survival in multiple cancers, when expressed at lower levels in the context of hTERT overexpression. Cell lines were classified into high and low quartiles according to hTERT gene expression.

**Figure S4: (A)** Validation of the SDL hits in the GTelo+/GTelo isogenic model that expresses Cas9 represented as a heatmap. The relative abundance of hTERT^high^ cells compared to hTERT^low^ cells is represented as the shade of green. Avg.PD is the average percentage difference in abundance between GTelo+ and GTelo-cells. **(B)** Validation results of the SDL hits in GTelo-/GTelo+ cells are presented. Briefly, equal numbers of Cas9-expressing isogenic cells were seeded in 96-well plates and subsequently transduced with lentiviruses expressing individual sgRNAs targeting each of the SDL hits or control sgRNA. The plates were imaged for 7 days by S3-IncuCyte®, and the cells were quantified to assess SDL effects. Multiple t-tests, *p < 0.01. **(C)** qRT‒PCR was used to measure and reconfirm hTERT levels in cell lines that were selected based on hTERT expression data available in Cancer Cell Line Encyclopedia. **(D)** Western blot showing Cas9 levels in two different cell lines for each tissue type selected for validation.

**Figure S5: (A)** Confirmation of genome editing by CRISPR/Cas9 in GTelo-Cas9 cells for several of the validated SDL hits are presented as examples. The cells were transduced with lentiviral particles containing two sgRNAs targeting individual hits, after which a cleavage assay was performed. The cleaved bands, representing genome editing, are shown in the lanes with endonuclease treatment (E+). **(B)** Expansion of the indicated cells that naturally overexpress hTERT was monitored using an S3-IncuCyte®. The red lines indicate knockouts of the indicated SDL genes of interest, and the blue lines indicate matching nontargeting controls. The asterisk represents at least one significant drop in confluency in one or more time points, calculated via multiple t-tests, *p < 0.01.

**Figure S6: (A)** Bar graph representing the qPCR data showing FTSJ3 gene expression values relative to the corresponding GAPDH levels. Primary pancreatic cancer cells are in green, hTERT^positive^ cells are in blue, ALT cells are in red, and non-malignant cells are in gray. **(B)** Bar graph depicting FTSJ3 knockdown efficiency relative to shRFP controls after normalization to GAPDH levels. **(C)** Western blot demonstrating decreased protein levels of FTSJ3 when transduced with shFTSJ3.

**Figure S7: (A)** Representative images depicting heterochromatin aggregates at the perinucleolus in hTERT^positive^ FTSJ3 knockdown cells. **(B)** Representative images of acrocentric centromere probes in shRFP and shFTSJ3 conditions. **(C)** Representative images of zoning and binning of telomeric signals into respective nuclear locations computationally via MetaExpress.

**Figure S8: (A)** Representative immunofluorescence of MIA PaCa-2 cells transduced with shRFP or shFTSJ3. Nucleus: DAPI; nucleolin; green; H3K9Me3 (top panel), H4Ac (bottom panel); red. N: nucleolus. Arrows indicate heterochromatin aggregates. **(B)** Subtelomeric (STH) H3K9me3 trimethylation (blue) and AcH4 acetylation (red) profiles of MIA PaCa-2 and U-2 OS for all chromosomes. For each chromosome, the rows (from top to bottom) represent subtelomeric H3K9me3, shRFP; subtelomeric H3K9me3, shFTSJ3; subtelomeric AcH4, shRFP; subtelomeric AcH4, shFTSJ3. For the subtelomeric plots, the most distal 5% regions of both arms are shown; these regions are depicted in the chromosome illustrations above each section. The x-axes show the base position in Mbp, and the y-axes show the methylation score. The data represent the mean methylation and acetylation values across two biological replicates (N=2). Peaks were smoothed with a Savitzky–Golay filter using a 100 bp window and a first-degree polynomial.

**Figure S9.** Pericentromeric (PCH) H3K9me3 trimethylation (blue) and AcH4 acetylation (red) profiles of MIA PaCa-2 and U-2 OS for all chromosomes. For each chromosome, the rows (from top to bottom) represent pericentromeric H3K9me3, shRFP; pericentromeric H3K9me3, shFTSJ3; pericentromeric AcH4, shRFP; and pericentromeric AcH4, shFTSJ3. For the pericentromeric plots, the regions comprising 5% of each chromosome from the canonical edge of the centromere are shown; these regions are depicted in the chromosome illustrations above each section. The x-axes show the base position in Mbp, and the y-axes show the methylation score. The data represent the mean methylation and acetylation values across two biological replicates (N=2). Peaks were smoothed with a Savitzky–Golay filter using a 100 bp window and a first-degree polynomial.

**Figure S10: (A)** Western blot demonstrating the autophosphorylation of ATM following FTSJ3 knockdown in hTERT^positive^ cells and ALT cells. **(B)** Western blot demonstrating no changes in p53, p21, 53BP1, pCHK1, and γ-H2AX protein expression in MIA PaCa-2 with FTSJ3 loss. **(C)** Interactions of FTSJ3 were downloaded from the STRING database and presented as a network to depict some of the key interactors. **(D)** Primer extension-based assay (with high and low dNTP concentrations) used to measure 2’-O methylation of hTERC is presented. Two different concentrations of dNTPs were tested. For low concentrations (L), 0.01 mM dNTPs were used, and for high concentrations (H), 0.4 mM dNTPs were used. The red arrow indicates a band that appeared only at a low concentration (0.01 mM) of dNTPs, which is indicative of a 2’-O-methylated site.

**Figure S11: (A)** Western blot testing FTSJ3 and DDX5 presence in total lysates as well as in samples after immunoprecipitation using anti-DDX5. **(B)** hTERT-positive cell lines and known ALT cell lines for which gene essentiality data from CRISPR screens were available in DepMap were used to analyze the essentiality of FTSJ3, DDX21 and DDX5. Lower scores represent higher levels of essentiality. Non-parametric Wilcoxon rank-sum test, p < 0.05. **(C)** Primer extension-based assay (with high and low dNTP concentrations) for measuring 2’-O methylation of TERRA in MIA PaCa-2 cells day 2, 5, 8 and 13 post-selection following transduction. The red arrow indicates a band that appeared only at a low concentration (0.01 mM) of dNTPs, which is indicative of a 2’-O-methylated site. The bottom panel shows an enhanced region of interest of the 2’-O-methylated site for clarity. **(D)** A visual schematic of preparation of TERRA sequencing via Oxford Nanopore Technologies’ MinION platform.

**Figure S12: (A)** Immunofluorescence images demonstrating no change in SUZ12 expression or localization after knockdown of FTSJ3 in either MIA PaCa-2 cells. Western blot showing that the expression of SUZ12 and TRF2 was unaffected by FTSJ3 knockdown in both MIA PaCa-2 and U-2 OS cells. **(B)** (Left) Representative immunofluorescence of MIA PaCa-2 cells transduced with shRFP or shFTSJ3. Nucleus: DAPI; SMARCA5: red. Arrows indicate CLAs. The nucleolus is marked by a broken line. (Right) Western blot analysis measuring TRF2 and SUZ12 levels in MIA PaCa-2 and U-2 OS cells five days following shFTSJ3 transduction. **(C)** Total telomere signal intensity and number of telomere signals in MIA PaCa-2-Cas9 cells transduced with sgFTSJ3. Telomere positions were identified using a threshold, followed by a calculation of the center of gravity and integrated intensity for each telomere in arbitrary units based on CCCTAA sequence/probe intensity. The integrated intensity reflects telomere length. The total number of telomere signals represents the sum of telomeres detected in each cell population. Least squares method, **p < 0.01 **(D)** Total telomere signal intensity and number of telomere signals in MIA PaCa-2 and U-2 OS cells transduced with shFTSJ3. Telomere positions were identified using a threshold, followed by calculation of the center of gravity and integrated intensity for each telomere in arbitrary units, based on TCA sequence/probe intensity. The integrated intensity reflects telomere length. The total number of telomere signals represents the sum of telomeres detected in each cell population. Chi-square, *p < 0.05. **(E)** FTSJ3 was silenced in U-2 OS cells, and dot blot analysis was subsequently performed to quantify the DNA:RNA hybrids. Equal amounts of DNA were spotted onto nitrocellulose membranes, and R-loops were detected using the S9.6 antibody. RNaseH treatment was also conducted, and the resulting product was used as a negative control to confirm the specificity of the signal. The ssDNA signal was used as an internal sample loading control.

**Figure S13: (A)** Transcriptome analysis of genes expression changes following FTSJ3 knockdown with greater than 50% change, p < 0.05. **(B)** (Top) Go-biological process enrichment of upregulated genes identified by RNAseq in MIA PaCa-2 following FTSJ3 knockdown (p < 0.05). (Middle) Go-biological process enrichment of upregulated genes identified by RNAseq in U-2 OS following FTSJ3 knockdown. (Bottom) Go-biological process enrichment of genes downregulated in RNAseq analysis of MIA PaCa-2 following FTSJ3 knockdown (p < 0.05). **(C)** The statistical significance of differences between the volumes of knockdown and control (shRFP) xenograft tumors was evaluated by an unpaired heteroscedastic t-test at the indicated time points following the inoculation of cancer cells. **(D)** The survival plot illustrates the difference in survival between patients with high TERT activity and low FTSJ3 activity (blue, n = 1,521) and the remaining patients (red, n = 3,340) within the selected category. Kaplan-Meier analysis yielded a log-rank p-value of 0.003082.

## Supplementary Tables

**Supplementary Table S1**: Significant hits identified in the four genome-wide pooled shRNA and CRISPR screens.

**Supplementary Table S2**: Reactome Pathways Enrichment Analysis

**Supplementary Table S3**: Interactions from IPA & screen.

**Supplementary Table S4**: Genes selected for validation

**Supplementary Table S5**: ALT and TERT cell lines used for analysis of DepMap gene essentiality scores.

**Supplementary Table S6**: Cell lines used in this work with its culture medium

**Supplementary Table S7**: Primers list for genome editing assay and CRISPR / microarray library sequencing.

