## Supplemental Figures for "Epigenetic Control of TERRA by FTSJ3 is Critical for Telomerase-Driven Cancers"

Supp Fig S1

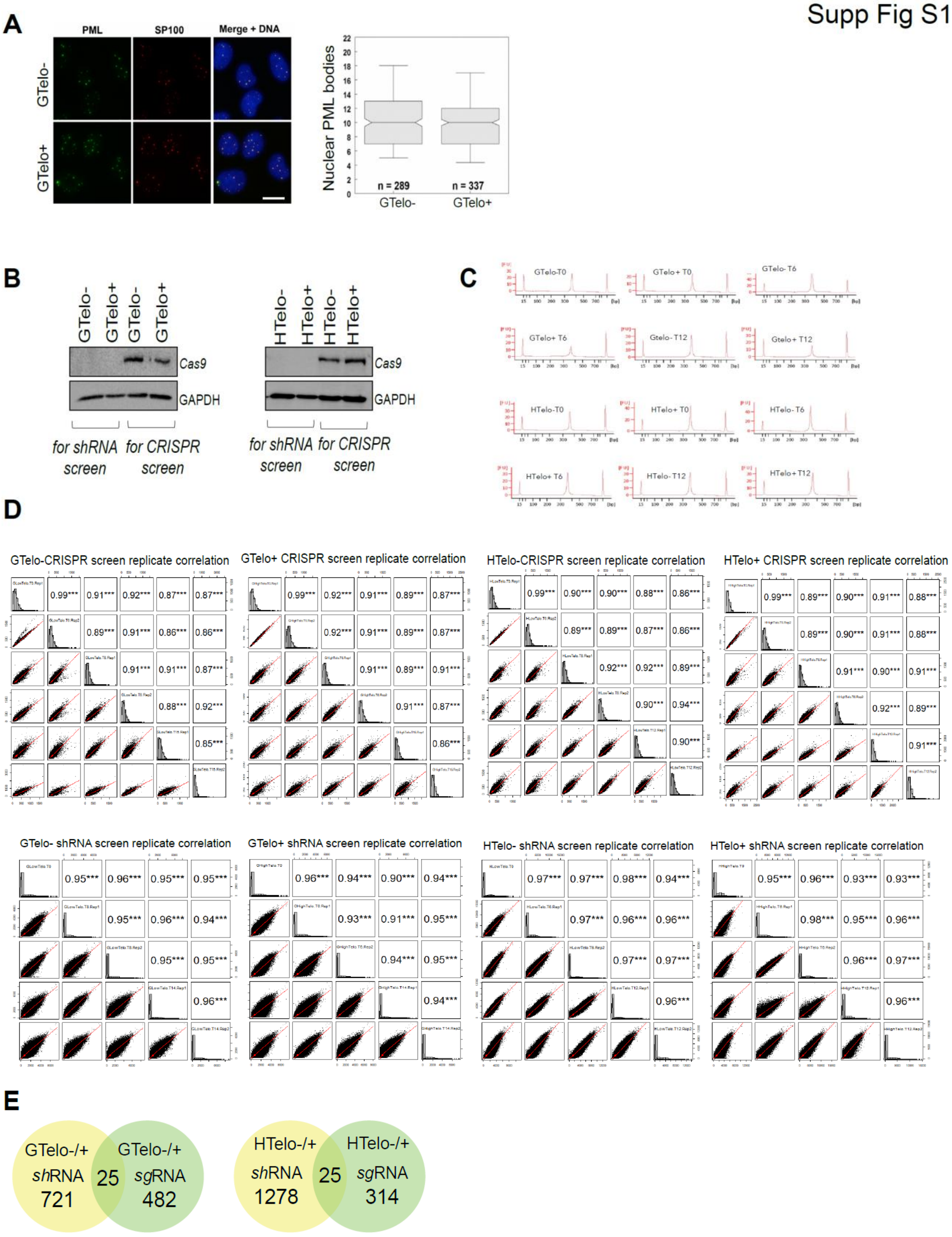
A
PML
SP100
Merge + DNA
GTelo-
GTelo+
Nuclear PML bodies
n = 289
n = 337
GTelo-
GTelo+
B
GTelo-
GTelo+
GTelo-
GTelo+
Cas9
GAPDH
for shRNA screen
for CRISPR screen
HTelo-
HTelo+
HTelo-
HTelo+
Cas9
GAPDH
for shRNA screen
for CRISPR screen
C
GTelo-T0
GTelo+ T0
GTelo- T6
GTelo+ T6
Gtelo- T12
Gtelo+ T12
HTelo-T0
HTelo+ T0
HTelo- T6
HTelo+ T6
HTelo- T12
HTelo+ T12
D
GTelo-CRISPR screen replicate correlation
GTelo+ CRISPR screen replicate correlation
HTelo-CRISPR screen replicate correlation
HTelo+ CRISPR screen replicate correlation
GTelo- shRNA screen replicate correlation
GTelo+ shRNA screen replicate correlation
HTelo- shRNA screen replicate correlation
HTelo+ shRNA screen replicate correlation
E
GTelo-/+ shRNA 721
25
GTelo-/+ sgRNA 482
HTelo-/+ shRNA 1278
25
HTelo-/+ sgRNA 314

Supp Fig S2

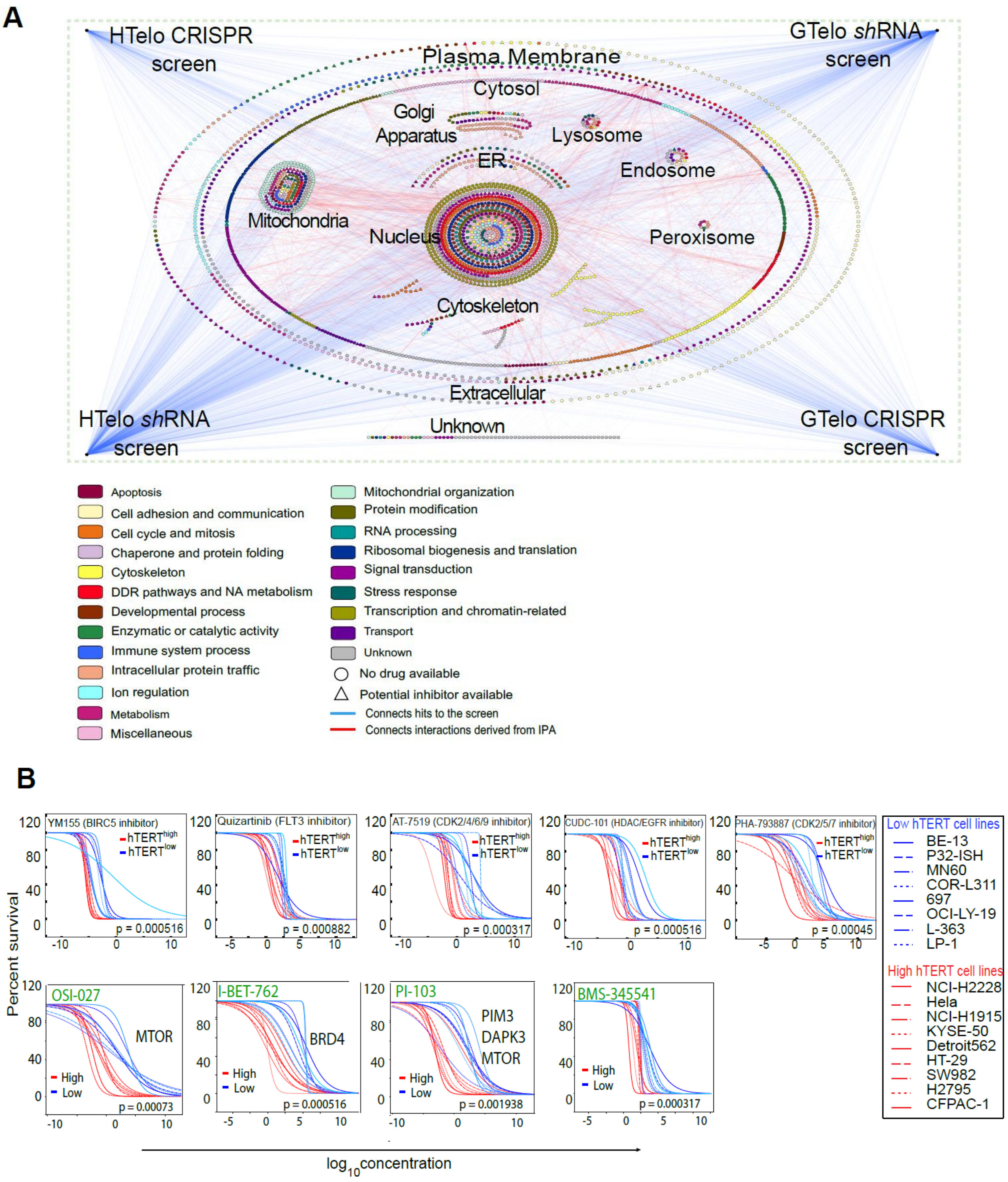
A
HTelo CRISPR screen
GTelo shRNA screen
HTelo shRNA screen
GTelo CRISPR screen
Plasma Membrane
Cytosol
Golgi Apparatus
Lysosome
ER
Endosome
Mitochondria
Nucleus
Peroxisome
Cytoskeleton
Extracellular
Unknown
Apoptosis
Cell adhesion and communication
Cell cycle and mitosis
Chaperone and protein folding
Cytoskeleton
DDR pathways and NA metabolism
Developmental process
Enzymatic or catalytic activity
Immune system process
Intracellular protein traffic
Ion regulation
Metabolism
Miscellaneous
Mitochondrial organization
Protein modification
RNA processing
Ribosomal biogenesis and translation
Signal transduction
Stress response
Transcription and chromatin-related
Transport
Unknown
No drug available
Potential inhibitor available
Connects hits to the screen
Connects interactions derived from IPA
B
YM155 (BIRC5 inhibitor)
p = 0.000516
Quizartinib (FLT3 inhibitor)
p = 0.000882
AT-7519 (CDK2/4/6/9 inhibitor)
p = 0.000317
CUDC-101 (HDAC/EGFR inhibitor)
p = 0.000516
PHA-793887 (CDK2/5/7 inhibitor)
p = 0.00045
hTERT^high
hTERT^low
OSI-027
MTOR
p = 0.00073
I-BET-762
BRD4
p = 0.000516
PI-103
PIM3
DAPK3
MTOR
p = 0.001938
BMS-345541
p = 0.000317
High
Low
Percent survival
log10concentration
Low hTERT cell lines
BE-13
P32-ISH
MN60
COR-L311
697
OCI-LY-19
L-363
LP-1
High hTERT cell lines
NCI-H2228
Hela
NCI-H1915
KYSE-50
Detroit562
HT-29
SW982
H2795
CFPAC-1

Supp Fig S3

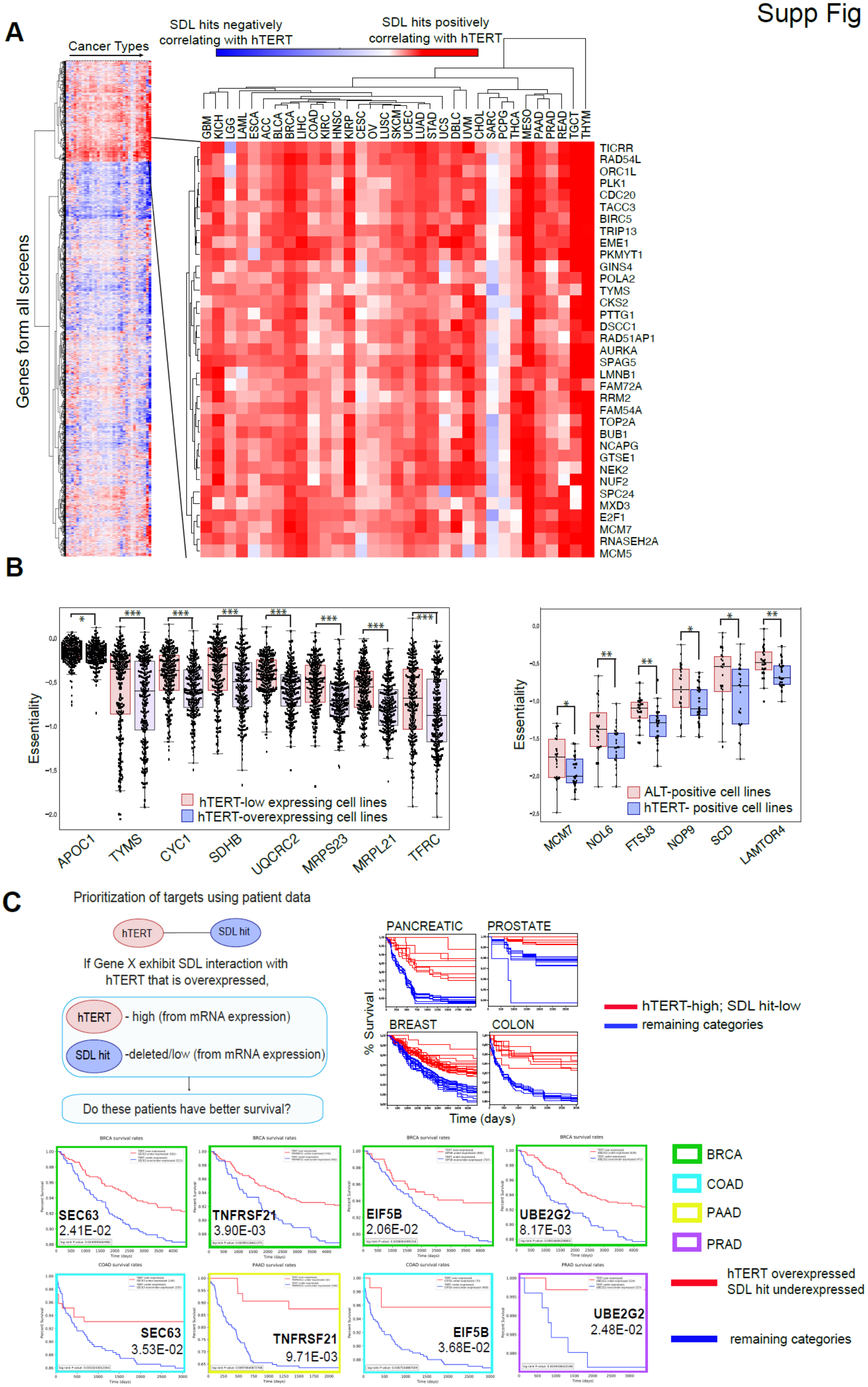
A
Cancer Types
SDL hits negatively correlating with hTERT
SDL hits positively correlating with hTERT
Genes form all screens
GBM
KICH
LGG
LAML
ESCA
ACC
BLCA
BRCA
LIHC
COAD
KIRC
HNSC
KIRP
CESC
OV
LUSC
SKCM
UCEC
LUAD
STAD
UCS
DBLC
UVM
CHOL
SARC
PCPG
THCA
MESO
PAAD
PRAD
READ
TGCT
THYM
TICRR
RAD54L
ORC1L
PLK1
CDC20
TACC3
BIRC5
TRIP13
EME1
PKMYT1
GINS4
POLA2
TYMS
CKS2
PTTG1
DSCC1
RAD51AP1
AURKA
SPAG5
LMNB1
FAM72A
RRM2
FAM54A
TOP2A
BUB1
NCAPG
GTSE1
NEK2
NUF2
SPC24
MXD3
E2F1
MCM7
RNASEH2A
MCM5
B
Essentiality
hTERT-low expressing cell lines
hTERT-overexpressing cell lines
APOC1
TYMS
CYC1
SDHB
UQCRC2
MRPS23
MRPL21
TFRC
ALT-positive cell lines
hTERT- positive cell lines
MCM7
NOL6
FTSJ3
NOP9
SCD
LAMTOR4
C
Prioritization of targets using patient data
hTERT
SDL hit
If Gene X exhibit SDL interaction with hTERT that is overexpressed,
hTERT - high (from mRNA expression)
SDL hit -deleted/low (from mRNA expression)
Do these patients have better survival?
PANCREATIC
PROSTATE
BREAST
COLON
% Survival
Time (days)
hTERT-high; SDL hit-low
remaining categories
BRCA survival rates
SEC63
2.41E-02
TNFRSF21
3.90E-03
EIF5B
2.06E-02
UBE2G2
8.17E-03
COAD survival rates
SEC63
3.53E-02
PAAD survival rates
TNFRSF21
9.71E-03
EIF5B
3.68E-02
PRAD survival rates
UBE2G2
2.48E-02
BRCA
COAD
PAAD
PRAD
hTERT overexpressed
SDL hit underexpressed
remaining categories

Supp Fig S4

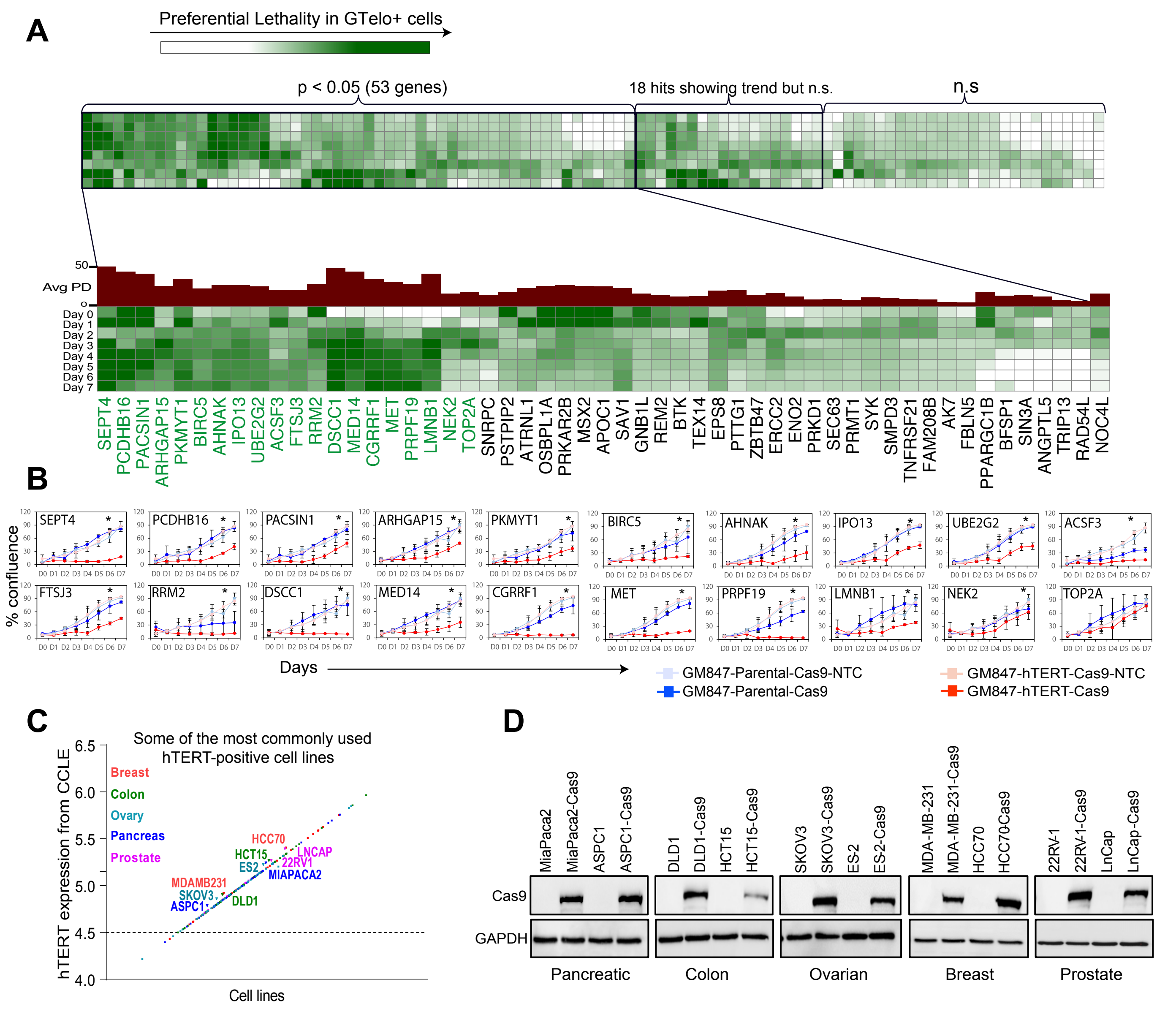
A
Preferential Lethality in GTelo+ cells
p < 0.05 (53 genes)
18 hits showing trend but n.s.
n.s
Avg PD
Day 0
Day 1
Day 2
Day 3
Day 4
Day 5
Day 6
Day 7
SEPT4 PCDHB16 PACSIN1 ARHGAP15 PKMYT1 BIRC5 AHNAK IPO13 UBE2G2 ACSF3 FTSJ3 RRM2 DSCC1 MED14 CGRRF1 MET PRPF19 LMNB1 NEK2 TOP2A SNRPC PSTPIP2 ATRNL1 OSBPL1A PRKAR2B MSX2 APOC1 SAV1 GNB1L REM2 BTK TEX14 EPS8 PTTG1 ZBTB47 ERCC2 ENO2 PRKD1 SEC63 PRMT1 SYK SMPD3 TNFRSF21 FAM208B AK7 FBLN5 PPARGC1B BFSP1 SIN3A ANGPTL5 TRIP13 RAD54L NOC4L
B
% confluence
SEPT4 PCDHB16 PACSIN1 ARHGAP15 PKMYT1 BIRC5 AHNAK IPO13 UBE2G2 ACSF3
FTSJ3 RRM2 DSCC1 MED14 CGRRF1 MET PRPF19 LMNB1 NEK2 TOP2A
Days
GM847-Parental-Cas9-NTC
GM847-Parental-Cas9
GM847-hTERT-Cas9-NTC
GM847-hTERT-Cas9
C
Some of the most commonly used hTERT-positive cell lines
hTERT expression from CCLE
Cell lines
Breast
Colon
Ovary
Pancreas
Prostate
HCC70
HCT15
LNCAP
ES2
22RV1
MIAPACA2
MDAMB231
SKOV3
ASPC1
DLD1
D
MiaPaca2 MiaPaca2-Cas9 ASPC1 ASPC1-Cas9 DLD1 DLD1-Cas9 HCT15 HCT15-Cas9 SKOV3 SKOV3-Cas9 ES-2 ES-2-Cas9 MDA-MB-231 MDA-MB-231-Cas9 HCC70 HCC70Cas9 22RV1 22RV-1-Cas9 LnCap LnCap-Cas9
Cas9
GAPDH
Pancreatic
Colon
Ovarian
Breast
Prostate

Supp Fig S5

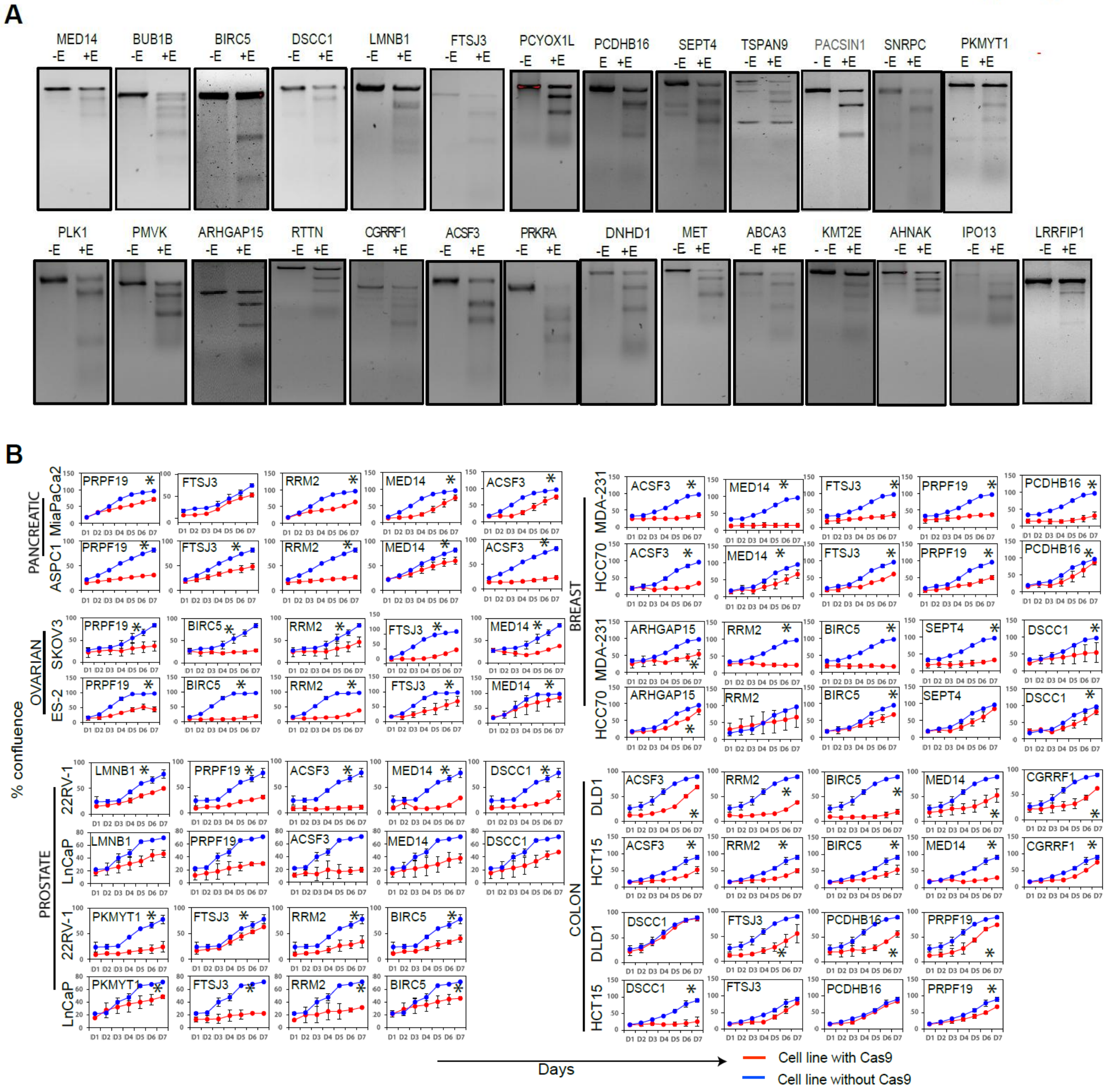
A
MED14 -E +E
BUB1B -E +E
BIRC5 -E +E
DSCC1 -E +E
LMNB1 -E +E
FTSJ3 -E +E
PCYOX1L -E +E
PCDHB16 E +E
SEPT4 -E +E
TSPAN9 -E +E
PACSIN1 - E +E
SNRPC -E +E
PKMYT1 E +E
PLK1 -E +E
PMVK -E +E
ARHGAP15 -E +E
RTTN -E +E
CGRRF1 -E +E
ACSF3 -E +E
PRKRA -E +E
DNHD1 -E +E
MET -E +E
ABCA3 -E +E
KMT2E - -E +E
AHNAK -E +E
IPO13 -E +E
LRRFIP1 -E +E
B
PANCREATIC
MiaPaCa2
ASPC1
OVARIAN
SKOV3
ES-2
PROSTATE
22RV-1
LnCaP
BREAST
MDA-231
HCC70
COLON
DLD1
HCT15
% confluence
Days
Cell line with Cas9
Cell line without Cas9

Supp Fig S6

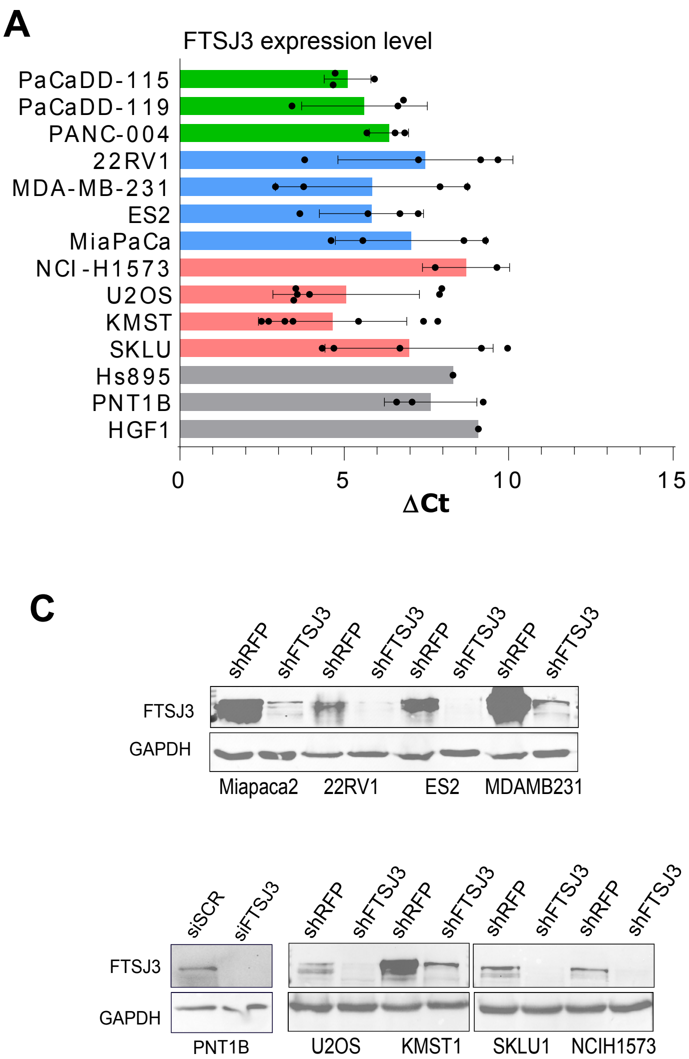

A
FTSJ3 expression level
PaCaDD-115
PaCaDD-119
PANC-004
22RV1
MDA-MB-231
ES2
MiaPaCa
NCI-H1573
U2OS
KMST
SKLU
Hs895
PNT1B
HGF1
0
5
10
15
ΔCt
B
FTSJ3 Knockdown efficiency
PaCaDD-115
PaCaDD-119
PANC-004
22RV1
MDA-MB-231
ES2
MiaPaCa2
NCI-H1573
U2OS
KMST1
SKLU1
Hs 895.SK
PNT1B
HGF-1
0.0 0.2 0.4 0.6 0.8 1.0 1.2
Fold gene expression (shRFP)
C
shRFP shFTSJ3 shRFP shFTSJ3 shRFP shFTSJ3 shRFP shFTSJ3
FTSJ3
GAPDH
Miapaca2 22RV1 ES2 MDAMB231
siSCR siFTSJ3 shRFP shFTSJ3 shRFP shFTSJ3 shRFP shFTSJ3 shRFP shFTSJ3
FTSJ3
GAPDH
PNT1B U2OS KMST1 SKLU1 NCIH1573

Supp Fig S7

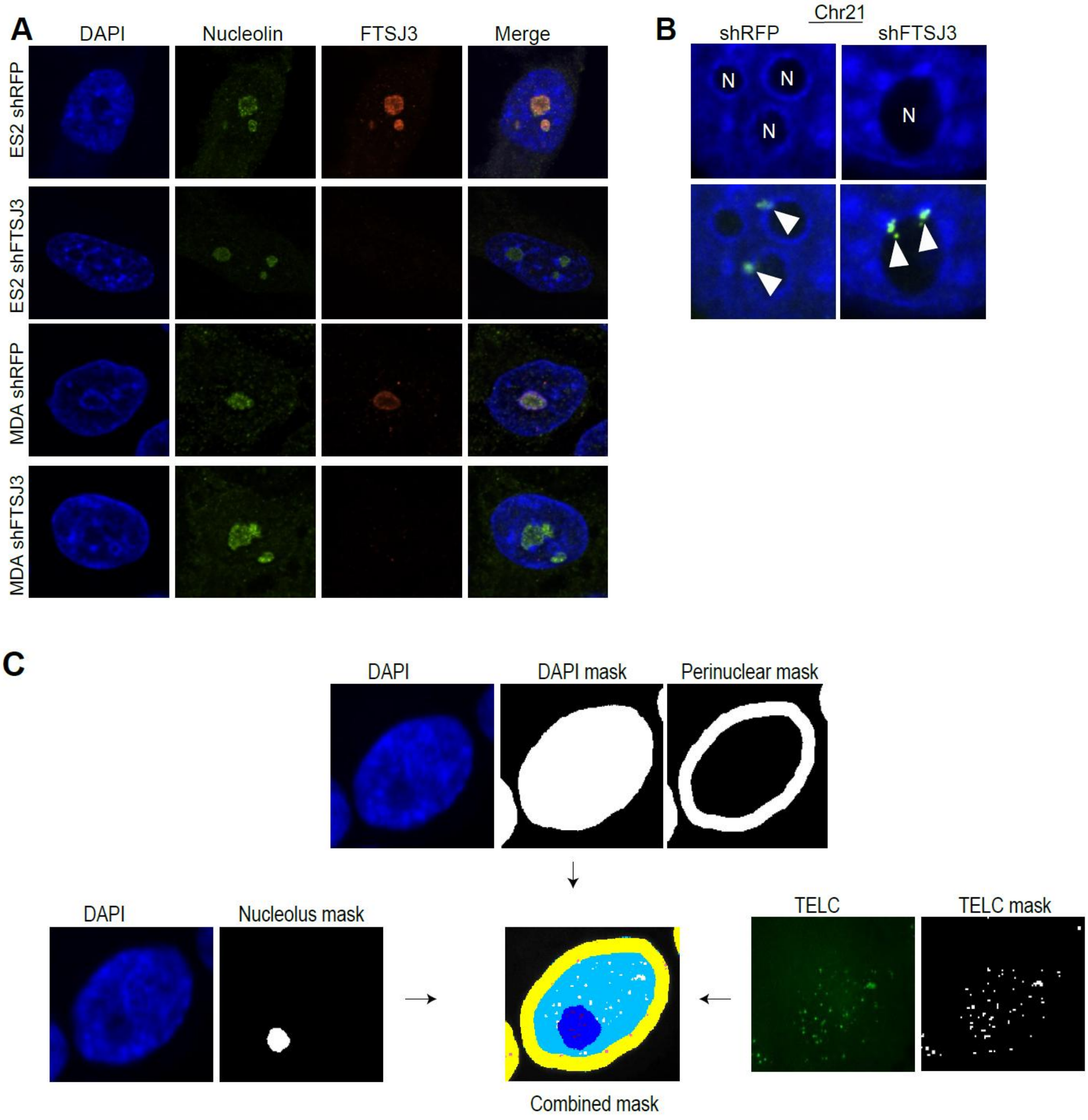
A
DAPI
Nucleolin
FTSJ3
Merge
ES2 shRFP
ES2 shFTSJ3
MDA shRFP
MDA shFTSJ3
B
Chr21
shRFP
shFTSJ3
N
N
N
N
C
DAPI
DAPI mask
Perinuclear mask
DAPI
Nucleolus mask
TELC
TELC mask
Combined mask

Supp Fig S8

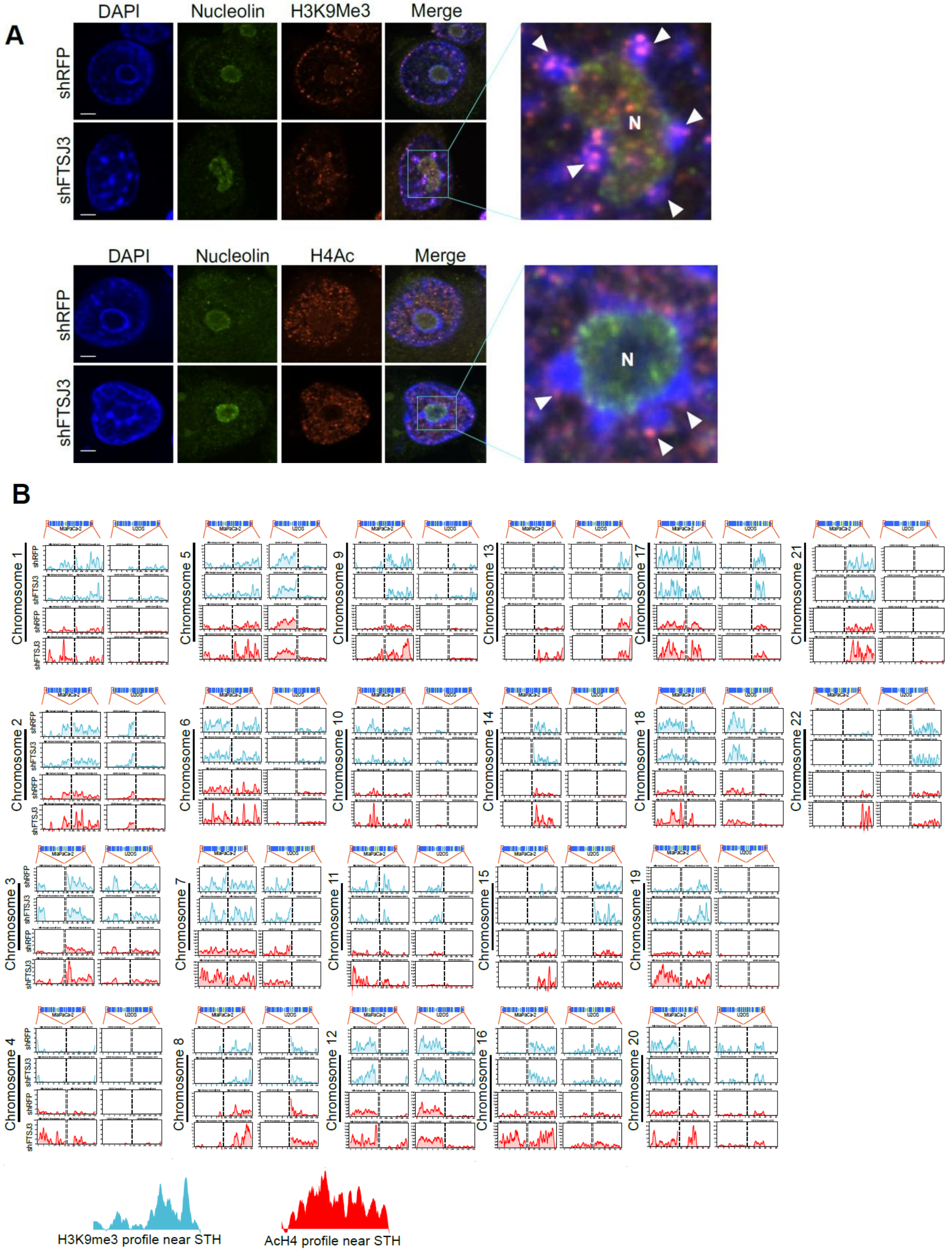
A
DAPI
Nucleolin
H3K9Me3
Merge
shRFP
shFTSJ3
N
DAPI
Nucleolin
H4Ac
Merge
shRFP
shFTSJ3
N
B
Chromosome 1
Chromosome 2
Chromosome 3
Chromosome 4
Chromosome 5
Chromosome 6
Chromosome 7
Chromosome 8
Chromosome 9
Chromosome 10
Chromosome 11
Chromosome 12
Chromosome 13
Chromosome 14
Chromosome 15
Chromosome 16
Chromosome 17
Chromosome 18
Chromosome 19
Chromosome 20
Chromosome 21
Chromosome 22
MiaPaCa-2
U2OS
shRFP
shFTSJ3
H3K9me3 profile near STH
AcH4 profile near STH

Supp Fig S9

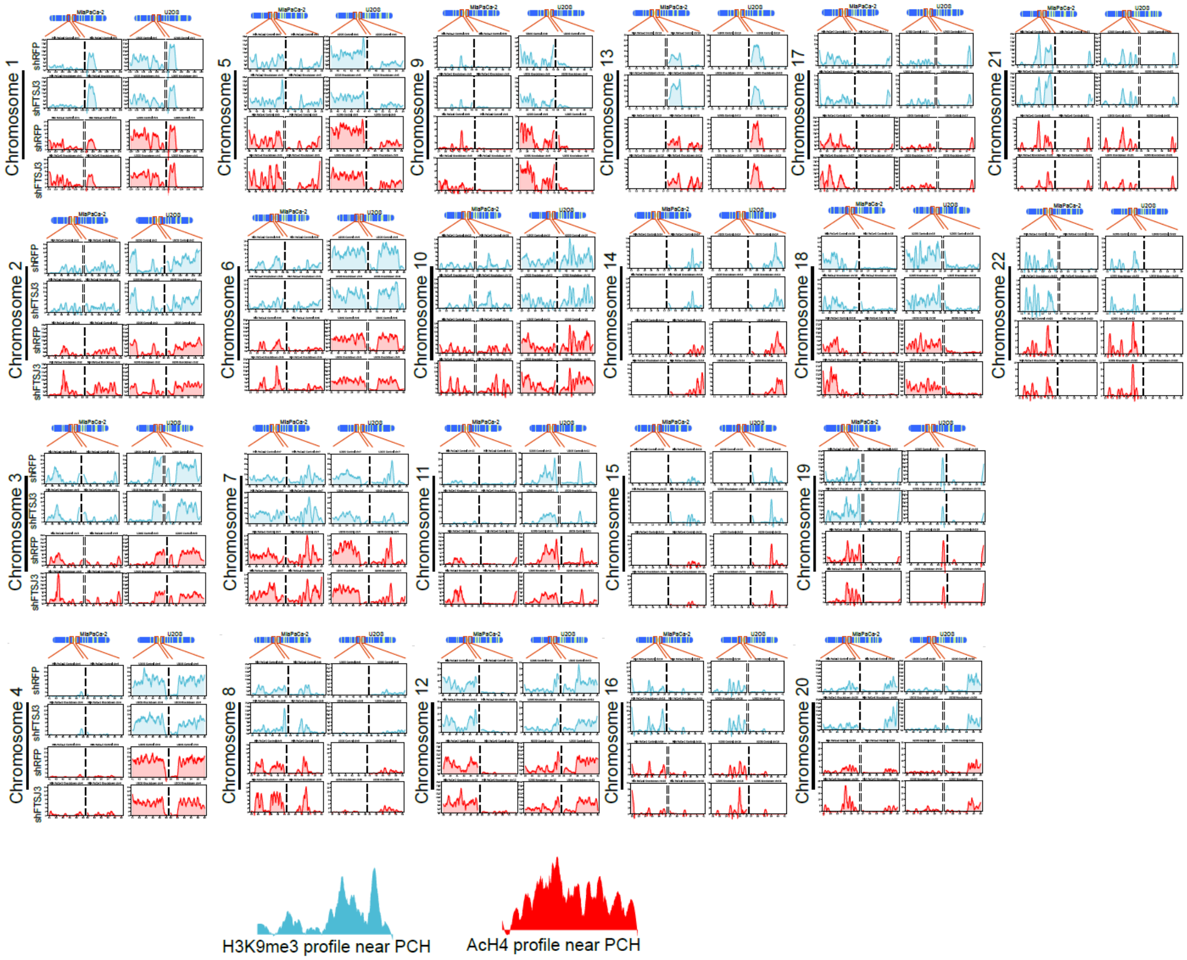
Chromosome 1
Chromosome 2
Chromosome 3
Chromosome 4
Chromosome 5
Chromosome 6
Chromosome 7
Chromosome 8
Chromosome 9
Chromosome 10
Chromosome 11
Chromosome 12
Chromosome 13
Chromosome 14
Chromosome 15
Chromosome 16
Chromosome 17
Chromosome 18
Chromosome 19
Chromosome 20
Chromosome 21
Chromosome 22
MiaPaCa-2
U2OS
shRFP
shFTSJ3
H3K9me3 profile near PCH
AcH4 profile near PCH

Supp Fig S10

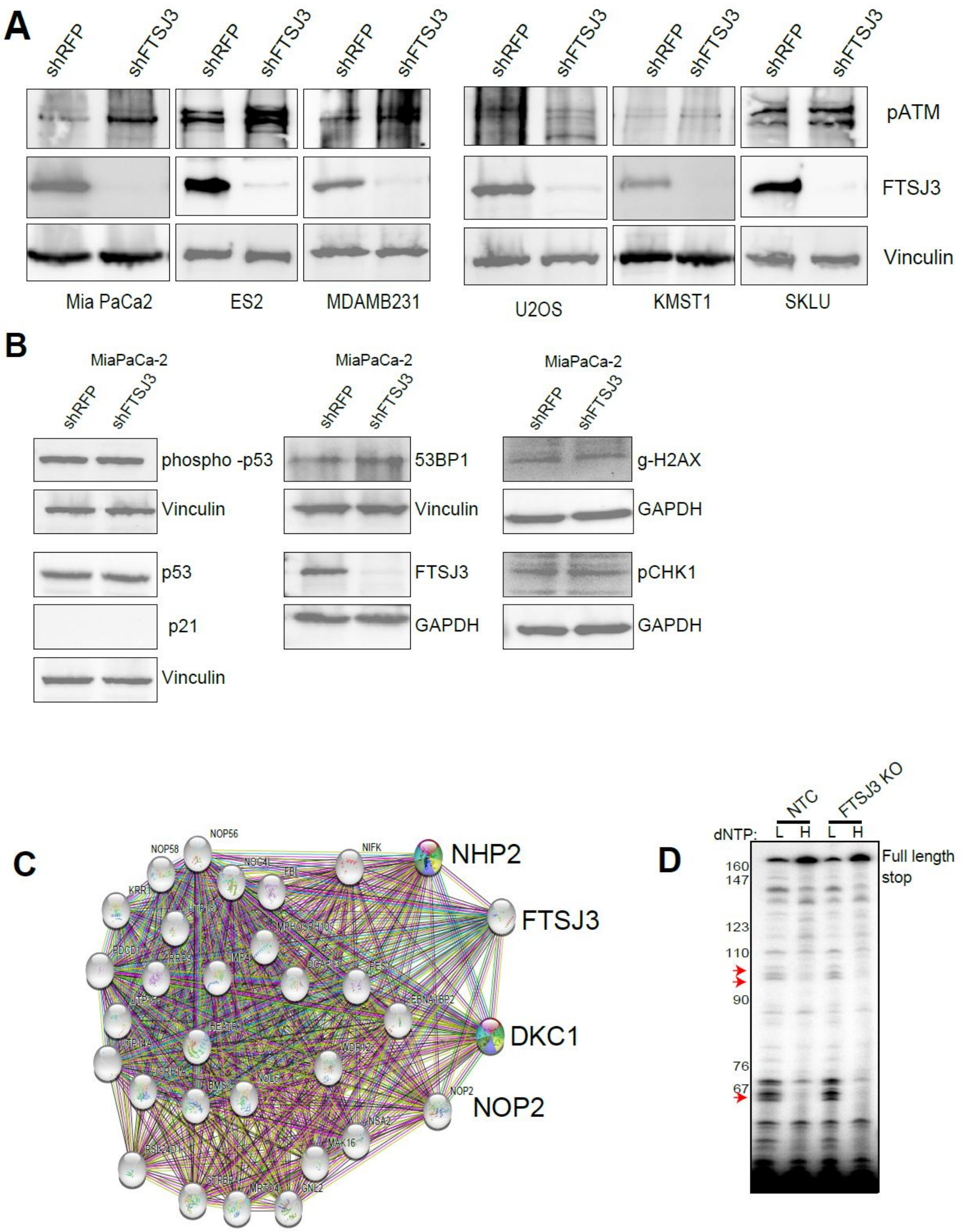
A
shRFP
shFTSJ3
pATM
FTSJ3
Vinculin
Mia PaCa2
ES2
MDAMB231
U2OS
KMST1
SKLU
B
MiaPaCa-2
phospho -p53
Vinculin
p53
p21
53BP1
FTSJ3
GAPDH
g-H2AX
pCHK1
C
NHP2
FTSJ3
DKC1
NOP2
NOP56
NOP58
NIFK
D
NTC
FTSJ3 KO
dNTP:
L
H
Full length
stop
160
147
123
110
90
76
67

Supp Fig S11

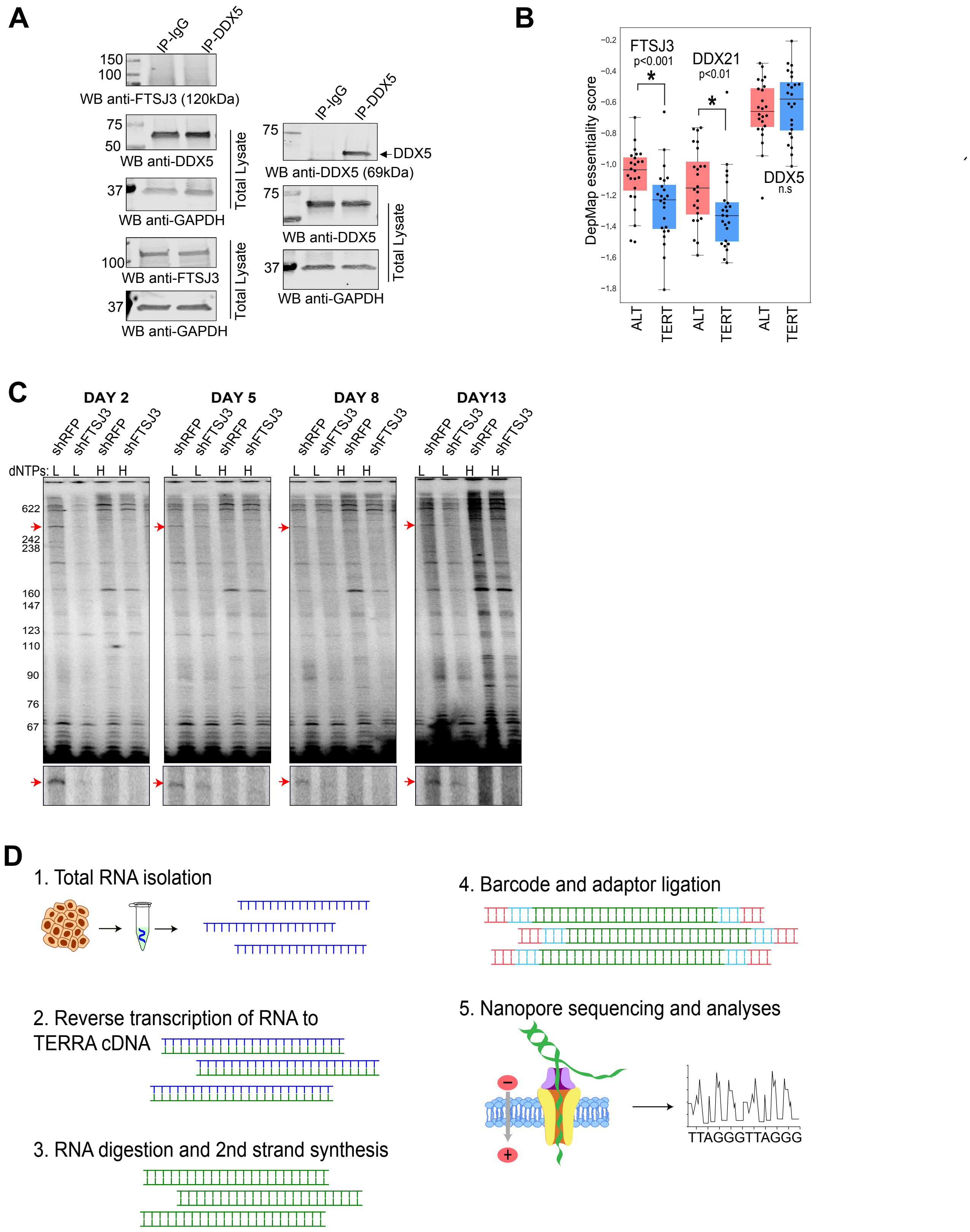
A
IP-IgG
IP-DDX5
150
100
WB anti-FTSJ3 (120kDa)
75
50
WB anti-DDX5
Total Lysate
37
WB anti-GAPDH
100
WB anti-FTSJ3
Total Lysate
37
WB anti-GAPDH
IP-IgG
IP-DDX5
75
DDX5
WB anti-DDX5 (69kDa)
75
WB anti-DDX5
Total Lysate
37
WB anti-GAPDH
B
FTSJ3
p<0.001
DDX21
p<0.01
DDX5
n.s
DepMap essentiality score
ALT
TERT
ALT
TERT
ALT
TERT
C
DAY 2
DAY 5
DAY 8
DAY13
shRFP
shFTSJ3
shRFP
shFTSJ3
dNTPs: L L H H
622
242
238
160
147
123
110
90
76
67
D
1. Total RNA isolation
2. Reverse transcription of RNA to TERRA cDNA
3. RNA digestion and 2nd strand synthesis
4. Barcode and adaptor ligation
5. Nanopore sequencing and analyses
TTAGGGTTAGGG

Supp Fig S12

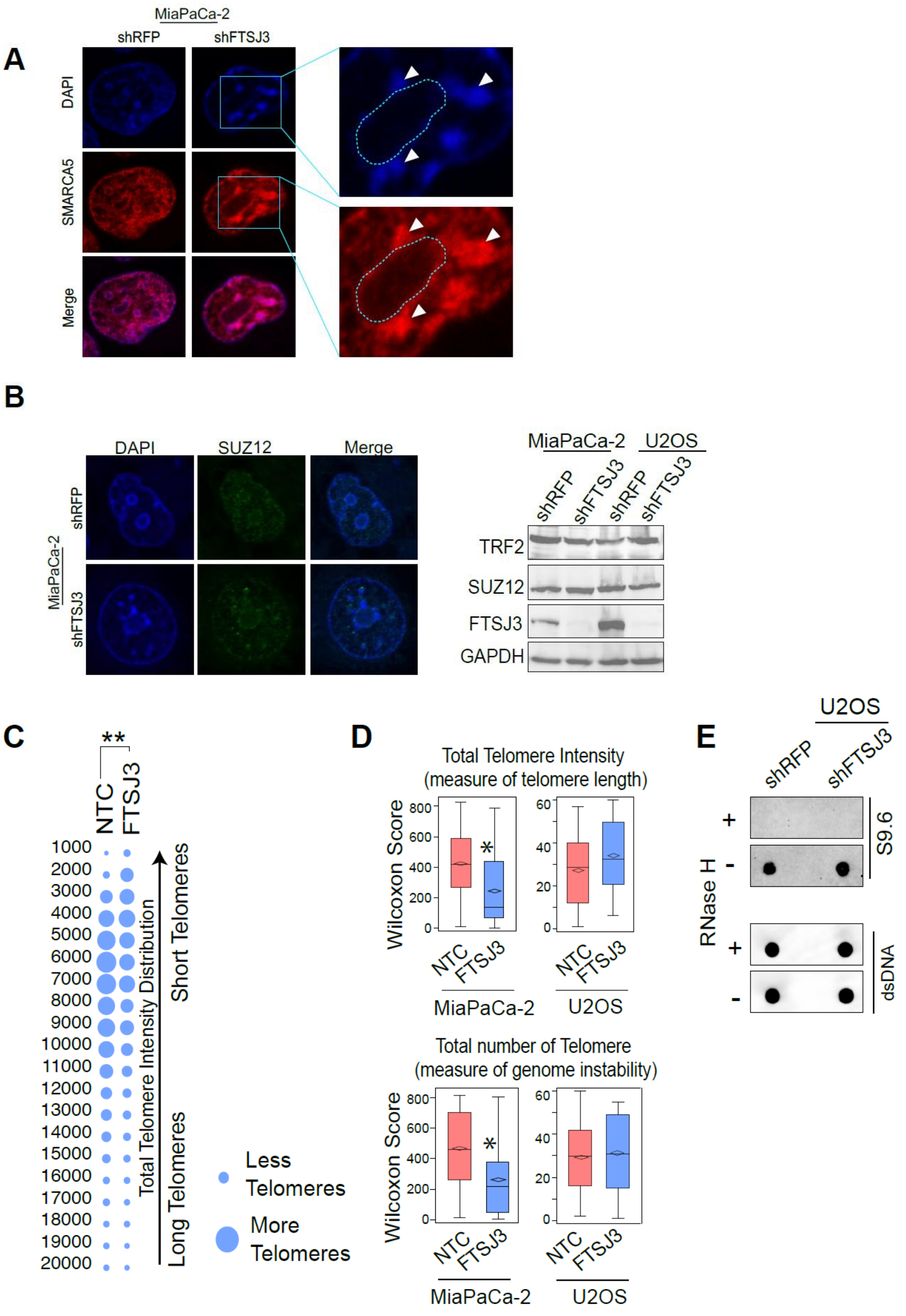

A
MiaPaCa-2
shRFP
shFTSJ3
DAPI
SMARCA5
Merge
B
DAPI
SUZ12
Merge
shRFP
MiaPaCa-2
shFTSJ3
MiaPaCa-2
U2OS
shRFP
shFTSJ3
shRFP
shFTSJ3
TRF2
SUZ12
FTSJ3
GAPDH
C
**
NTC
FTSJ3
1000
2000
3000
4000
5000
6000
7000
8000
9000
10000
11000
12000
13000
14000
15000
16000
17000
18000
19000
20000
Total Telomere Intensity Distribution
Short Telomeres
Long Telomeres
Less Telomeres
More Telomeres
D
Total Telomere Intensity
(measure of telomere length)
Wilcoxon Score
NTC
FTSJ3
MiaPaCa-2
U2OS
Total number of Telomere
(measure of genome instability)
Wilcoxon Score
NTC
FTSJ3
MiaPaCa-2
U2OS
E
U2OS
shRFP
shFTSJ3
RNase H
+
-
+
-
S9.6
dsDNA

Supp Fig S13

A

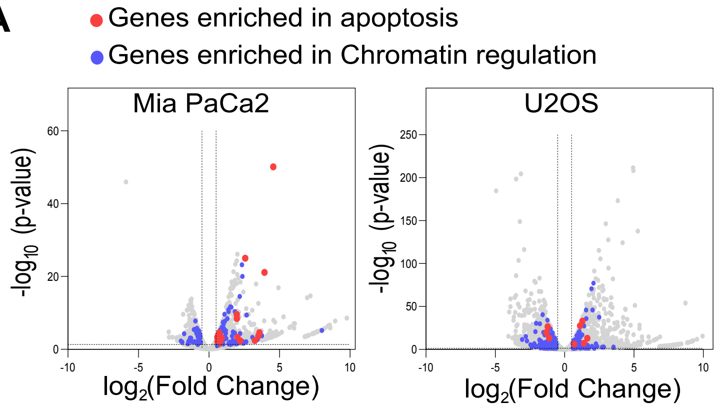


B

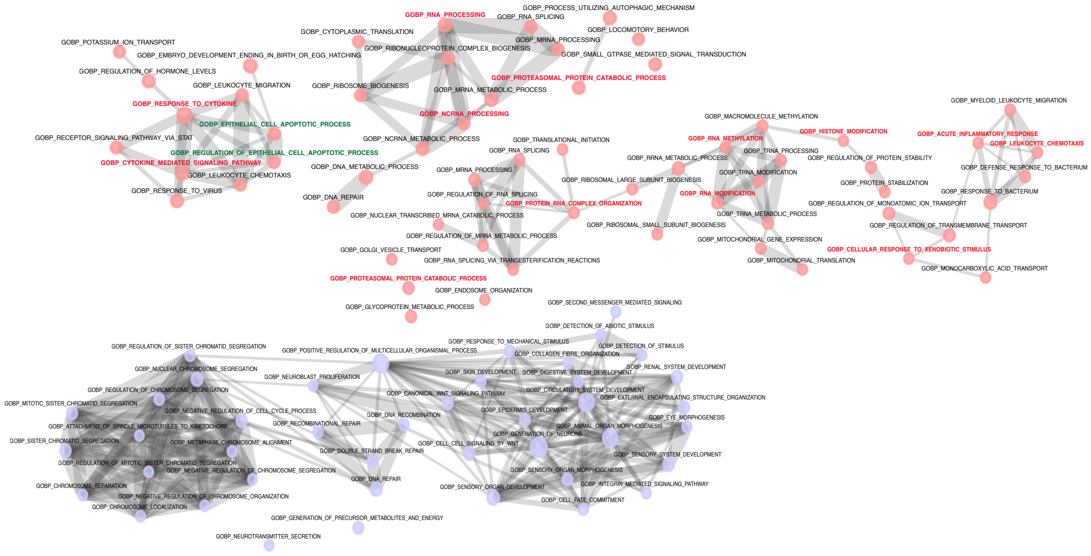

C

p-values for xenograft experiments

| 22Rv1 | Day 13 | Day 17 | Day 20 | Day 24 | Day27 | Day 31 | Day 34 | Day 38 |
|---|---|---|---|---|---|---|---|---|
| ***shFTSJ3*** | 0.1220 | 0.0060 | 0.0005 | 0.0109 | 0.0024 | 0.0008 | 0.0013 | 0.0015 |

| HCC70 | Day 21 | Day 25 | Day 28 | Day 31 | Day 35 |
|---|---|---|---|---|---|
| ***shFTSJ3*** | 0.0001 | 0.0004 | 0.0001 | 4.81E-05 | 2.21E-06 |

| ES2 | Day 11 | Day 14 | Day 18 | Day 21 |
|---|---|---|---|---|
| ***shFTSJ3*** | 0.0269 | 0.0024 | 0.0008 | 6.27E-07 |

| MiaPaca2 | Day 14 | Day 18 | Day 22 | Day 25 | Day 29 | Day 32 | Day 36 | Day 39 | Day 41 | Day 44 |
|---|---|---|---|---|---|---|---|---|---|---|
| ***shFTSJ3*** | 0.0935 | 0.1773 | 0.0303 | 0.0178 | 0.0041 | 0.0111 | 0.1202 | 0.2391 | 0.2549 | 0.3042 |

D

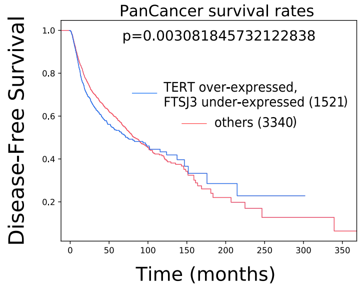
